# Microglia and Fn14 regulate transcription and chromatin accessibility in developing neurons

**DOI:** 10.1101/2021.08.16.456505

**Authors:** Austin Ferro, Uma Vrudhula, Yohan S.S. Auguste, Lucas Cheadle

**Affiliations:** Cold Spring Harbor Laboratory, Cold Spring Harbor, NY 11724

**Keywords:** Microglia, cytokine, sensory experience, synapse, refinement, Fn14, snRNAseq, ATACseq

## Abstract

Cytokine signaling pathways that promote inflammation in peripheral tissues are repurposed to coordinate the refinement of synaptic connections in the developing brain. However, the downstream mechanisms through which these pathways mediate neural circuit maturation remain to be fully defined. Here, we demonstrate that Fn14, a cytokine receptor that promotes inflammation outside of the central nervous system, shapes the transcriptional profiles and chromatin landscapes of neurons in the developing brain. Single-nucleus RNA-sequencing revealed hundreds of misregulated genes in the thalamocortical neurons of the visual thalami of mice lacking either Fn14 or its microglial derived cytokine ligand TWEAK, including genes encoding proteins with critical roles in synaptic function, histone modification, and chromatin remodeling. Whole-genome analysis uncovered significant alterations in chromatin accessibility in the brains of mice lacking Fn14 or in wild-type mice following microglial depletion, and chromatin changes due to both manipulations were enriched near genes encoding regulators of synaptic function. Loss of microglia also led to the retention of excess synapses, suggesting that microglia may link modifications in neuronal chromatin to the functional refinement of neural circuits. Consistent with Fn14 shaping brain function beyond the visual system, Fn14 knockout mice displayed impairments in memory task proficiency as well as heightened sensitivity to pharmacologically induced seizures. Taken together, these results define a previously undescribed interaction between microglia, cytokine signaling, and the neuronal epigenome that is likely to contribute to neural circuit refinement and function in the brain.

## Introduction

The precise organization of synaptic connections between defined neuronal partners is the basis of mature brain function. These connections are established in a step-wise fashion, beginning with the formation of an overabundance of synapses and concluding with a refinement process in which a subset of these synapses is strengthened and maintained while others are eliminated. While the initial assembly of synapses occurs largely through genetic mechanisms intrinsic to neurons, synaptic refinement is also coordinated extensively by sensory experience during critical periods in the postnatal brain (Hooks and Chen, 2020; Hubel and Wiesel, 1963; Wiesel and Hubel, 1963). These phases of heightened sensory-dependent (SD) refinement occur concurrently with the epigenomic maturation of neurons which is thought to contribute to the maintenance of these newly refined circuits in the long-term (Frank et al., 2015; Gallegos et al., 2018; Stroud et al., 2017; Stroud et al., 2020). Disruptions in both synaptic refinement and neuronal epigenomic maturation have been implicated in neurodevelopmental disorders such as autism and schizophrenia, and are also associated with the aberrant elimination of mature synapses in conditions such as Alzheimer’s disease (AD)(Feinberg, 1982; Hammond et al., 2019; LeBlanc and Fagiolini, 2011; Starr, 2019). However, the cellular and molecular mechanisms underlying the SD maturation of neurons and their synapses in the healthy brain remain unclear.

Over the past 15 years, the retinogeniculate circuit of the mouse has emerged as a leading model for studying synaptic refinement. In this circuit, retinal ganglion cells (RGCs) in the eye relay information about the visual world to downstream structures in the brain by projecting their axons onto excitatory thalamocortical (TC) neurons in the dorsal lateral geniculate nucleus (dLGN) of the thalamus. Electrophysiological analyses have shown that retinogeniculate refinement—i.e. the concurrent removal of some connections between RGCs and TC neurons and the strengthening of others—occurs across multiple distinct phases that are coordinated by different patterns of neural activity (Hong and Chen, 2011). During the first week of life, synaptic inputs from each eye that converge onto the same territory of the dLGN are removed, leading to the anatomical segregation of inputs based upon their eye of origin (eye-specific segregation, ESS) in a process that depends upon activity spontaneously generated in the retina (Corriveau et al., 1998; Shatz and Kirkwood, 1984). Once inputs are segregated, spontaneous retinal activity continues to drive the functional refinement of retinogeniculate synapses until P20. Finally, visual experience is required for the further refinement of synapses during a critical period that takes place between P20 and P30, after which the retinogeniculate circuit is thought to be largely mature (Hooks and Chen, 2006, 2008). Thus, the retinogeniculate circuit provides a robust experimental paradigm in which to study multiple aspects of neural circuit refinement.

A major contribution of the retinogeniculate circuit to our mechanistic understanding of synaptic refinement has been the discovery of roles for immune-related signaling pathways (e.g. cytokines and their receptors) and the brain’s resident immune cells, microglia, in neural circuit development. For example, groundbreaking work over the past decade has shown that microglia promote ESS during the first week of life by engulfing (or *phagocytosing*) RGC inputs through signaling between components of the classical complement cascade (ccc), a pathway that mediates innate immunity in the periphery. In brief, complement component 1q (C1q) and complement component 3 (C3) are secreted by microglia (and other cell types) and deposited onto a subset of immature synapses, thereby tagging them for removal. Recognition of synaptic C3 by the C3 Receptor (CR3) on microglia then elicits the elimination of the complement-tagged synapses through phagocytic engulfment (Schafer et al., 2012; Stevens et al., 2007). An extensive body of work emerging from these studies has gone on to define the roles of the ccc, and microglia in general, in synaptic refinement across numerous neural circuits and in the contexts of both health and disease (Hong et al., 2016; Lee et al., 2019). In parallel, a host of other immune-related signaling molecules—including fractalkine, interleukin-33, and C1q-like proteins—have been implicated in orchestrating refinement in the retinogeniculate circuit and beyond (Gunner et al., 2019; Kakegawa et al., 2015; Nguyen et al., 2020). Hence, the repurposing of cytokine pathways that mediate inflammation in peripheral tissues has emerged as a major mechanism underlying synaptic refinement in the developing brain.

The vast majority of cytokines known to be involved in neural circuit development function by eliciting the remodeling or phagocytosis of individual synapses through localized interactions between synapses and microglia. Whether microglia and cytokines also orchestrate aspects of circuit development beyond local changes at synapses—for example, the epigenomic maturation of neurons which occurs at the same time as synaptic refinement—has remained unclear. In addition, the roles of microglia and cytokine signaling specifically during the late phase of SD refinement which, in the dLGN, is characterized by markedly low levels of synaptic phagocytosis, are still being elucidated. In addressing these gaps in knowledge, we recently identified signaling between the microglia-derived Tumor Necrosis Factor (TNF) superfamily cytokine TNF-associated Weak inducer of apoptosis (TWEAK) and its neuronal receptor Fibroblast Growth Factor-inducible protein 14 kDa (Fn14) as essential for SD refinement in the dLGN (Cheadle et al., 2020; Cheadle et al., 2018). TWEAK and Fn14 mediate a diverse array of functions outside of the brain (e.g. skeletal muscle remodeling after injury, liver development, inflammation, angiogenesis, and cell migration) by inducing transcriptional programs in Fn14-expressing cells that encode key mediators of cellular remodeling (Burkly, 2014; Dogra et al., 2007; Jakubowski et al., 2005; Meighan-Mantha et al., 1999; Tran et al., 2006). This observation raised the possibility that TWEAK-Fn14 signaling may link microglia to SD synaptic refinement not through localized interactions between microglia and synapses, but through more global mechanisms involving the regulation of neuronal transcription. However, the possibility that microglia engage cytokine signaling to influence the transcriptomic and epigenomic profiles of developing neurons remained to be tested.

Given the precise involvement of TWEAK-Fn14 signaling in sensory experience-dependent refinement and the observation that this pathway eliminates synapses through a mechanism that is distinct from phagocytic engulfment, defining how TWEAK and Fn14 promote refinement could uncover new ways in which microglia mediate brain development and disease. In this study, we employed single-nucleus RNA-sequencing (snRNAseq) to characterize the gene targets of TWEAK and Fn14 in TC neurons of the dLGN, revealing the misregulation of genes involved in synaptic plasticity and organization, histone modification, and chromatin remodeling in the absence of either signaling partner. Consistent with Fn14 actively contributing to the regulation of transcription in neurons, chromatin accessibility profiling revealed a large number of non-coding *cis*-regulatory regions that exhibit differential accessibility in the absence of Fn14. A subset of these regions was predicted to regulate the expression of genes that we found to be misregulated in the Fn14 KO by snRNAseq. Depletion of microglia from the brain also led to significant alterations in the chromatin landscape of neurons while simultaneously impeding the SD elimination of synapses. Regulatory regions whose accessibility is mediated by Fn14 or microglia, though largely non-overlapping, were predicted to control the expression of genes involved in synaptic signaling and neurotransmission. Finally, we show that Fn14 is expressed across a variety of brain structures in adult mice, including the hippocampus, and that loss of Fn14 disrupts memory function and increases seizure susceptibility *in vivo*. Altogether, these results reveal that microglia and the cytokine receptor Fn14 regulate transcription and chromatin accessibility in developing neurons, and suggest that this process is critical for the development and function of the healthy brain.

## Results

### Whole-transcriptome characterization of Fn14-regulated genes at single-cell resolution

The ability of TWEAK and Fn14 to coordinate transcriptional changes in response to injury outside of the brain led us to hypothesize that TWEAK and Fn14 refine synapses at least in part by modifying gene expression in developing neurons. Given that Fn14 is the neuronal receptor of the TWEAK-Fn14 pathway and that Fn14 expression is restricted to TC neurons in the dLGN (Fig. 1A), we reasoned that if this pathway regulates neuronal transcription, we would observe robust transcriptional changes in the TC neurons of mice lacking Fn14. Thus, to identify the genes that are regulated by Fn14 in TC neurons in an unbiased manner, we performed single-nucleus RNA-sequencing (snRNAseq) on the micro-dissected dLGNs of Fn14 KO and WT littermates at P27, a time point at which we have shown Fn14 to promote synapse refinement in response to sensory experience (Cheadle et al., 2018). We utilized the inDrops method of high-throughput sequencing to sample the transcriptomes of individual nuclei across two biological replicates of each genotype (Zilionis et al., 2017). After the removal of putative doublets and sequencing read quality control (Fig. S1A-D), we proceeded with downstream analysis of a final dataset of 11,586 nuclei using the R package Seurat v3 (McGinnis et al., 2019; Stuart et al., 2019). Nuclei were assigned to with excitatory TC neuron clusters being identified by co-expression of *Slc17a7* (*Vglut1*), *Slc17a6* (*Vglut2*), and *Prkcd* (Fig. 1B,C and Fig. S2A-G)(Tasic et al., 2016). While the clustering algorithm separated TC neurons into two large and two very small clusters, their separation was based upon relatively subtle transcriptional differences, consistent with prior studies (Bakken et al., 2021; Kalish et al., 2018). Thus, we focused the bulk of our initial analysis on TC clusters one and two as these contained by far the largest numbers of TC cells, increasing our ability to detect differentially expressed genes across conditions. These clusters were sequenced to an average depth of 2985 and 1740 unique molecular identifiers (UMIs) per cell, respectively (see STAR Methods).

**Figure 1.**
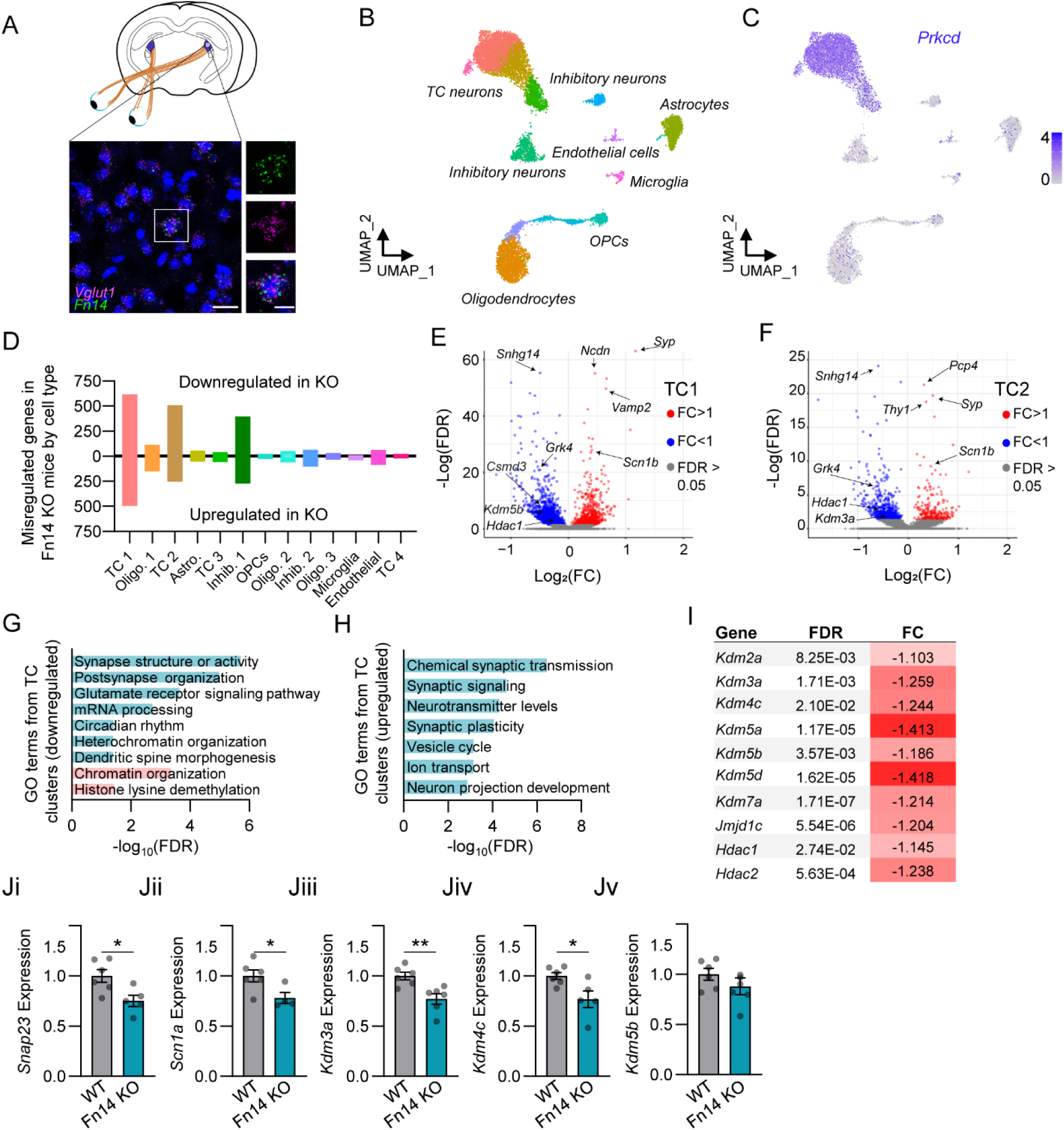
Characterization of Fn14-regulated genes in thalamocortical (TC) neurons of the dLGN by snRNAseq. (A) Fluorescence *in situ* hybridization of the thalamocortical marker *Vglut1* and *Fn14* in the dLGN (scale bars = 20 μm). (B) UMAP (Uniform Manifold Approximation and Projection for Dimension Reduction) plot displaying cell clusters present in the snRNAseq dataset. (C) Selective expression of the TC (thalamocortical) neuron marker *Prkcd* in TC neuron clusters (color intensity scale: log normalized transcripts per cell). (D) Histogram displaying the numbers of differentially expressed genes (FDR < 0.05, Log2FC > |±1.25|) between Fn14 WT and KO mice. Genes downregulated in the Fn14 knockout (KO), above x-axis; genes upregulated in the KO, below x-axis. See also figures S1 and S2, and supplemental tables 1, and 2. (E),(F) Volcano plots of differentially expressed genes in TC clusters 1 (E) and 2 (F). Genes significantly downregulated in the KO shown in blue, non-significantly changed genes in grey, and significantly upregulated genes in the KO shown in red. (G) GO (gene ontology, PANTHER terms) categories enriched in genes downregulated in Fn14 KO neurons. Teal bars, GO terms shared by TC clusters 1 and 2; pink bars, terms downregulated in TC cluster 1 specifically. (H) GO terms from shared upregulated genes in TC clusters 1 and 2. See also supplemental table 3. (I) Table of histone lysine demethylases (KDMs) and histone deacetylases (Hdacs) downregulated in Fn14 KO TC custer 1. (Ji-Jv) qPCR analysis of whole brain tissue validating the downregulation of select genes in Fn14 KO mice normalized to *GAPDH* expression. KO values normalized to WT values. Data represent the mean ± SEM with individual data points representing individual mice (n = 4-6 mice/group). Unpaired, 2-tailed student’s T test (*p < 0.05, **p < 0.01). Cell cluster abbreviations: Thalamocortical neurons (TC), Oligodendrocytes (Oligo.), Astrocytes (Astro.), Inhibitory neurons (Inhib.), Oligodendrocyte precursor cells (OPCs).

We next utilized the R package Monocle v2 to identify genes that were significantly differentially expressed (<u>d</u>ifferentially <u>e</u>xpressed <u>g</u>enes, DEGs) in WT versus Fn14 KO cells (Qiu et al., 2017). DEG analysis of TC clusters one and two revealed that gene expression is robustly misregulated in neurons from the Fn14 KO mouse. In Fn14 KO cells of TC cluster one, 616 genes were downregulated and 479 genes were upregulated compared to WT at a false discovery rate of less than 5% (FDR < 0.05) and a fold-change threshold of greater than 1.25-fold (Fig. 1D, E)(supplemental table 1). In the second TC cluster, 510 genes were downregulated while 237 genes were upregulated in Fn14 KO neurons (Fig. 1D, F)(supplemental table 2). Thus, loss of Fn14 led to a bidirectional misregulation of genes with a modest majority of genes (56% in cluster 1 and 68% in cluster 2) being downregulated rather than upregulated in the Fn14 KO. Consistent with these transcriptional changes arising from a defined role for Fn14 in mediating transcription rather than as an indirect result of the relative circuit immaturity previously characterized in the Fn14 KO mouse, the majority of DEGs in the dataset were observed in TC neurons specifically, which are by far the most prominent expressers of Fn14 in the dLGN (Fig. 1A). While we also noted substantial transcriptional changes in an inhibitory neuron cluster distinguished by high levels of Proenkephalin (*Penk)* expression, fluorescence *in situ* hybridization (FISH) confirmed that these cells reside in nearby structures outside of the dLGN, and that a small number of these cells was inadvertently captured in our dLGN microdissection (Fig. S3). Thus, the vast majority of transcriptional changes in the dLGNs of Fn14 KO mice were observed in TC neurons, as expected.

We next performed gene ontology analyses to assess the functions of Fn14- regulated genes broadly in an unbiased manner (Mi et al., 2013). Consistent with Fn14 mediating synapse elimination by controlling neuronal transcription, a substantial number of misregulated genes encoded molecules with important functions at synapses. Downregulated genes in Fn14 KO TC clusters included genes associated with glutamate receptor signaling (e.g. glutamate receptors *Gria2* and *Grik1*), dendritic spine morphogenesis (e.g. the membrane trafficking protein *Dnm3*), and postsynaptic organization (e.g. cell adhesion molecules *Lrrtm2* and *NrCAM*)(Fig. 1G, teal)(supplemental table 3). Among the molecules encoded by genes in these functional classes, the cytoskeletal regulator Kalirin is a promising candidate effector of SD refinement given its roles in the activity-dependent plasticity of dendritic spines and the observation that its expression in the developing brain peaks at P28 (Xie et al., 2007).

Another gene that stands out as a potential effector of synapse elimination is the cytoskeletal regulator Dock7, mutations in which contribute to epilepsy and deficits in visual function (Perrault et al., 2014; Tai et al., 2014). Similar to the genes that were downregulated in the Fn14 KO mouse, those that were upregulated also encoded proteins with synaptic functions. However, whereas downregulated genes were more closely associated with the postsynaptic domain, genes that were upregulated in the Fn14 KO also included molecules with important roles in presynaptic neurotransmission (Fig. 1H). For example, TC neurons lacking Fn14 displayed heightened expression of genes involved in synaptic vesicle cycling (e.g. vesicular associated membrane proteins *Vamp1* and *Vamp2,* as well as *Synaptophysin*) and neuron projection development (e.g. the cation channel *Scn1b* and the cytoskeletal remodelers *Rac1* and *Pak1*) (Fig 1E, H). Given that Fn14 is required for synapse elimination, we speculate that the decreased expression of these genes in the presence of Fn14 may tune the molecular composition of synapses to favor their disassembly. Overall, loss of Fn14 led to the bidirectional misregulation of critical synaptic organizers in TC neurons.

While a substantial proportion of Fn14-regulated genes are directly involved in the remodeling of synapses, a separate cohort is specialized to regulate more global aspects of neuronal maturation. Specifically, we observed that critical mediators of histone modification and chromatin organization are also significantly downregulated (but not upregulated) in the TC neurons of Fn14 KO mice (Fig. 1E-G). The proteins encoded by downregulated genes include both Hdac1 and -2, enzymes that repress transcriptional activation by removing acetylation marks from histones, and Dnmt3a, a protein that methylates DNA in a postnatal process that establishes stable gene expression as the brain matures (Park and Kim, 2020; Stroud et al., 2017). In addition, a relatively large number of histone lysine demethylases (KDMs)—including (KDM)2a, -3a, -4c, -5a, -5b, - 5d, -7a, and Jmjd1c—were found to be downregulated in Fn14 KO neurons (Fig. 1G, I, J). This observation is particularly noteworthy given known roles for KDMs in brain development and their strong association with human neurodevelopmental disorders (Hatch et al., 2021; Hyun et al., 2017; Wijayatunge et al., 2014; Wijayatunge et al., 2018). In line with Fn14 functioning in part by regulating gene expression through varied mechanisms beyond histone modification, genes downregulated in Fn14 KO neurons were also enriched for mediators of mRNA splice site selection, RNA 3’ end processing, and RNA export from the nucleus (Fig. 1G). Thus, Fn14 likely exerts control over the transcriptomic maturation of neurons through the genomic regulation of gene transcription and through the regulation of transcribed RNA. Altogether, these data reveal an important role for Fn14 in the expression of genes encoding local synaptic proteins as well as regulators of the chromatin landscapes of TC neurons in the dLGN. Given the critical role for Fn14 in SD synapse refinement (Cheadle et al., 2018), these data uncover Fn14 as a potential link between circuit development and the epigenomic maturation of neurons in the brain.

### The microglial cytokine TWEAK regulates gene expression in the dLGN

While Fn14 can regulate gene expression in the absence of its ligand TWEAK, TWEAK requires Fn14 to mediate its cellular functions (Brown et al., 2013; Dogra et al., 2007). Therefore, we hypothesized that genetic deletion of TWEAK would result in the misregulation of gene expression in dLGN TC neurons as we observed in the Fn14 KO mouse. To identify TWEAK-regulated genes, we performed snRNAseq on the dLGNs of TWEAK KO and WT mice across four bioreplicates per genotype using the same approaches as described above. The final dataset included 50,004 nuclei of which we focused the bulk of our analysis on TC neuron cluster two because it exhibited the highest read-depth of TC neuron clusters at 2025.62 UMIs per cell (Fig. 2A,B; Fig. S1E-H; Fig. S2H-O). Identification of DEGs across all cell types using Monocle v2 revealed that, unlike in the Fn14 KO mouse, the changes in gene expression were not primarily observed in TC cells but distributed much more evenly across the diversity of cell types in the dataset (Fig. 2C). Nevertheless, in TC cluster 2, we found that 68 genes were upregulated and 109 genes were downregulated in the absence of TWEAK (Fig. 2C, E)(supplemental table 4). Consistent with Fn14 mediating transcription and synaptic refinement both in the presence and absence of TWEAK, the overall numbers of misregulated genes in TC neurons of the TWEAK KO were much lower than in the Fn14 KO, which was particularly striking for the genes that are downregulated in the respective KOs (Fig. 2Di). Conversely, the numbers of genes that were upregulated in Fn14 and TWEAK KO mice were more closely aligned (Fig. 2Dii).

**Figure 2.**
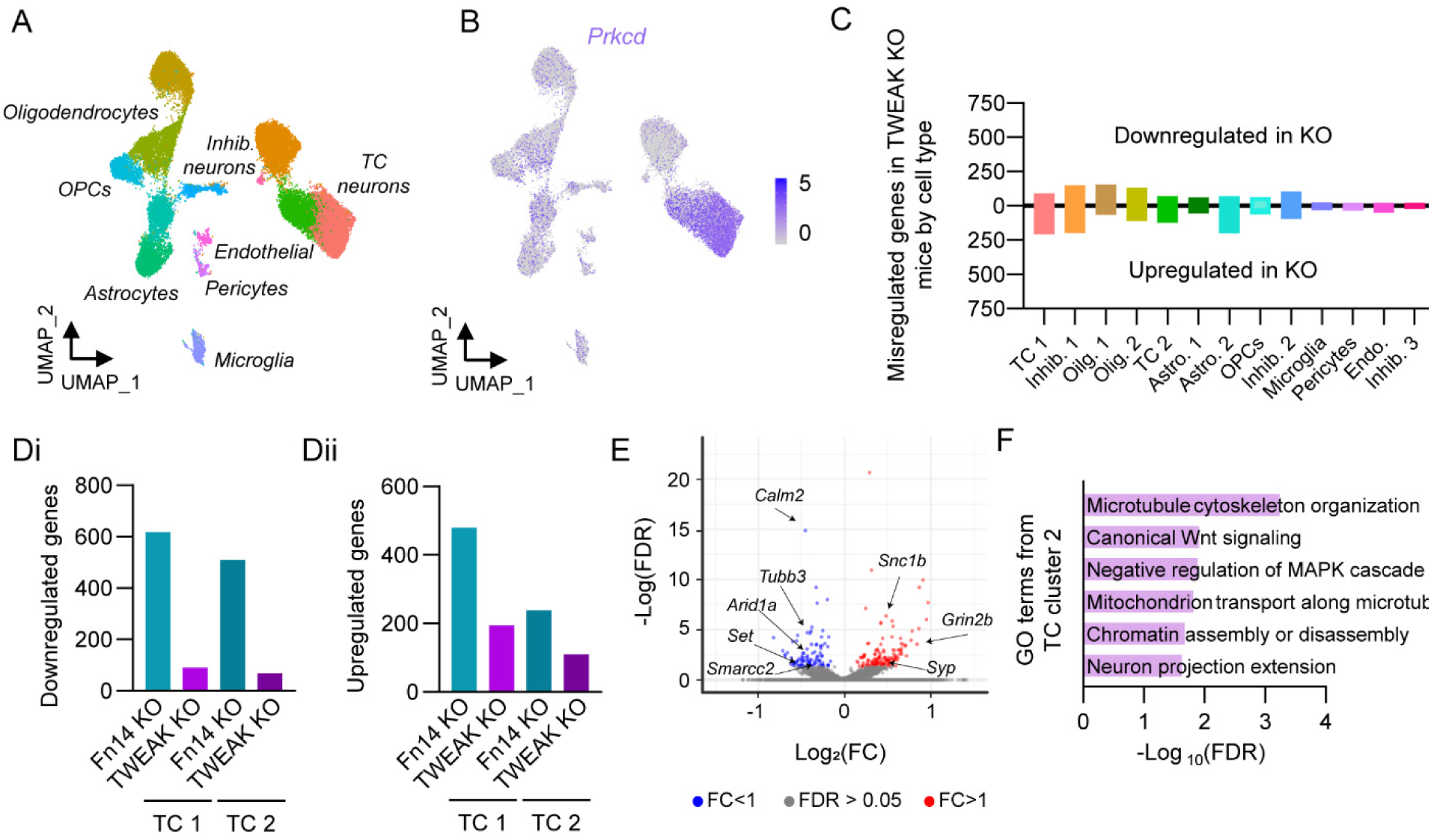
Characterization of TWEAK-regulated genes in thalamocortical (TC) neurons of the dLGN by snRNAseq. (A) UMAP (Uniform Manifold Approximation and Projection for Dimension Reduction) plot displaying cell clusters present in the snRNAseq dataset. (B) Selective expression of the TC neuron marker *Prkcd* in TC neuron clusters (color intensity scale, log normalized transcripts per cell). (C) Histogram displaying the numbers of differentially expressed genes (FDR < 0.05, Log2FC > |±1.25|) between TWEAK WT and knockout (KO) mice. Genes downregulated in the KO, above x-axis; genes upregulated in the KO, below x-axis. Y-axis scaled for comparison with graph in Fig. 1D. See also figures S1 and S2. (Di) Comparison of the numbers of genes significantly downregulated in Fn14 and TWEAK KO mice, TC clusters 1 and 2. (Dii) Comparison of the numbers of genes significantly upregulated in Fn14 and TWEAK KO mice, TC clusters 1 and 2. (E) Volcano plot of differentially expressed genes in TC cluster 2. Genes downregulated in the TWEAK KO shown in blue, non-significantly changed genes in grey, and genes upregulated in the TWEAK KO shown in red. (F) Significantly enriched GO terms from downregulated genes in TWEAK KO cluster 2. See also supplemental tables 4 and 5. Cell cluster abbreviations: Thalamocortical neurons (TC) Oligodendrocytes (Oligo.), Astrocytes (Astro.), Inhibitory neurons (Inhib.), Oligodendrocyte precursor cells (OPCs), Endothelial cells (Endo.).

Comparison of the misregulated genes in the Fn14 and TWEAK KO mice with one another revealed a low degree of overlap between these gene sets. This finding is likely to reflect that the misregulated genes that we detect in the Fn14 KO neurons are enriched for targets whose expression does not require TWEAK, but could also be the result of the constitutive loss of TWEAK expression leading to compensatory mechanisms limiting our ability to detect the full complement of TWEAK targets. Nevertheless, some of the TWEAK-regulated genes that we identified encode mediators of the same biological processes as the genes that are regulated by Fn14, while others are predicted to function in distinct aspects of circuit development. For example, mediators of postsynaptic development and function were not as prevalent among the TWEAK-regulated genes as among Fn14-regulated genes, whereas TWEAK-regulated genes were enriched for putative mediators of presynaptic development such as neuronal process extension and microtubule organization (e.g. *Map1b* and several members of the Tubulin family)(supplemental table 5).

The most intriguing observation to arise from this analysis was the enrichment of regulators of chromatin assembly and disassembly among TWEAK-regulated genes, emphasizing that TWEAK and Fn14 may both couple SD refinement to long-term changes in the chromatin landscape (Fig. 2F). For example, TWEAK regulates the expression of *Set*, an enzyme that inhibits the acetylation of histones (Bannister and Kouzarides, 2011; Bryk et al., 2002; Kim et al., 2013). TWEAK also regulates the expression of two members of the SWI/SNF nuclear complex of chromatin remodelers, *Smarcc2* and *Arid1a*, which mediate the function of enhancers (non-coding regions of DNA that potentiate or repress the transcription of associated genes) through various mechanisms. Mutations in the genes encoding these remodelers are associated with neurodevelopmental disorders such as autism (Alver et al., 2017; Bogershausen and Wollnik, 2018). Overall, these data suggest that the TWEAK-Fn14 pathway mediates SD refinement not only by coordinating the expression of direct regulators of synapse development, but also by inducing histone modifications and chromatin changes to facilitate the maturation of dLGN TC neurons as their connectivity is refined. The data also suggest that separate cohorts of genes are regulated by Fn14 through self-association versus through TWEAK-Fn14 binding.

### Transcriptional misregulation in non-neuronal cells of TWEAK and Fn14 KO mice

While we focused the bulk of our snRNAseq analysis on TC neurons as the cellular locus of TWEAK-Fn14 signaling, an advantage of the snRNAseq approach is that it also allowed us to assess the more global effects of removing TWEAK and Fn14 from the developing brain. Thus, we also analyzed the misregulation of gene expression in non-neuronal cell classes of the dLGN. We observed a particularly large number of misregulated genes in oligodendrocytes from both TWEAK and Fn14 KO mice, including the upregulation of genes involved in axon development such as the myelin associated protein *Mag*. Similarly, astrocytes from both TWEAK and Fn14 KO mice displayed decreased expression of genes involved in ion homeostasis and neurotransmitter uptake (e.g. *Kcnj10* and *Slc6a1*), consistent with their roles in facilitating neurotransmission. Fn14 KO astrocytes were also found to misregulate Alzheimer’s disease associated genes *ApoE* and *APP*, an interesting observation given the proposed association between TWEAK-Fn14 signaling and neurodegeneration (Nagy et al., 2021). Finally, while microglia were minimally affected by loss of Fn14, TWEAK KO microglia upregulated genes involved in neurotransmitter reuptake such as *Slc1a2*, which may represent a compensatory mechanism to limit excitotoxicity given the aberrant retention of excess synapses in these mice (Cheadle et al., 2020)(Fig. S4). Taken together, these data suggest that TWEAK-Fn14 signaling modestly, but significantly, impacts non-neuronal transcription, likely through non-cell-autonomous pathways initiated by changes in the expression of cell:cell signaling molecules in neurons. Overall, the snRNAseq analyses in both TWEAK and Fn14 KO mouse lines uncovered a genome-wide pattern of transcriptional misregulation when TWEAK-Fn14 signaling, or TWEAK-independent Fn14 function, is disrupted.

### Loss of Fn14 alters the chromatin landscape of neurons in the developing brain

A major function of histone modification and chromatin remodeling is to shape the genomic landscape such that specific *cis*-regulatory regions of the genome become accessible (or inaccessible) to binding by transcription factors (TFs), nuclear proteins that can activate, potentiate, or repress transcription. Dynamic changes in chromatin accessibility therefore have the potential to determine which genes are (or are not) expressed and the degree of their expression. Given that Fn14 and TWEAK regulate the expression of a variety of chromatin and histone modifying enzymes (e.g. Hdacs, KDMs, and members of the SWI/SNF complex) predicted to have diverse effects on gene regulation, we next sought to identify how the decreased expression of these factors (via genetic loss of Fn14) affects chromatin accessibility in developing neurons. To this end, we mapped chromatin accessibility across the genome by performing <u>A</u>ssay for <u>T</u>ransposase-<u>A</u>ccessible <u>C</u>hromatin followed by <u>seq</u>uencing (ATACseq) in the brains of Fn14 KO and WT mice at P27 (Buenrostro et al., 2015)(Fig. S5A). Following alignment and quality-filtering using the Encode Pipeline (https://www.encodeproject.org/atac-seq/), we ran the DiffBind package to identify significantly differentially accessible peaks between conditions from a set of 194,231 total consensus peaks identified across all samples. The most differentially accessible peak that we uncovered was associated with the Fn14 promoter and this peak was absent in the Fn14 KO, validating the approach (Fig. S5B). Remarkably, we identified 700 total regions of the genome that are significantly more or less accessible upon loss of Fn14, with 197 sites being less accessible and 503 sites being more accessible in the Fn14 KO (FDR < 0.05)(Fig. 3A-C,E,F)(supplemental table 6). Analysis of the differentially accessible peaks using EnrichR validated that, although our dissection included multiple posterior brain regions, the samples were enriched for excitatory neurons, the predominant expressers of Fn14 (Fig. S5Di)(Xie et al., 2021). The vast majority of the chromatin accessibility changes in the Fn14 KO mouse represented at least a 2-fold difference in accessibility between the genotypes, and there was strong agreement regarding accessibility across both bioreplicates (Fig. S5E,F). Thus, Fn14 mediates the accessibility of a defined cohort of *cis*-regulatory regions across the genome, consistent with the misregulation of chromatin-organizing genes in the Fn14 KO.

**Figure 3.**
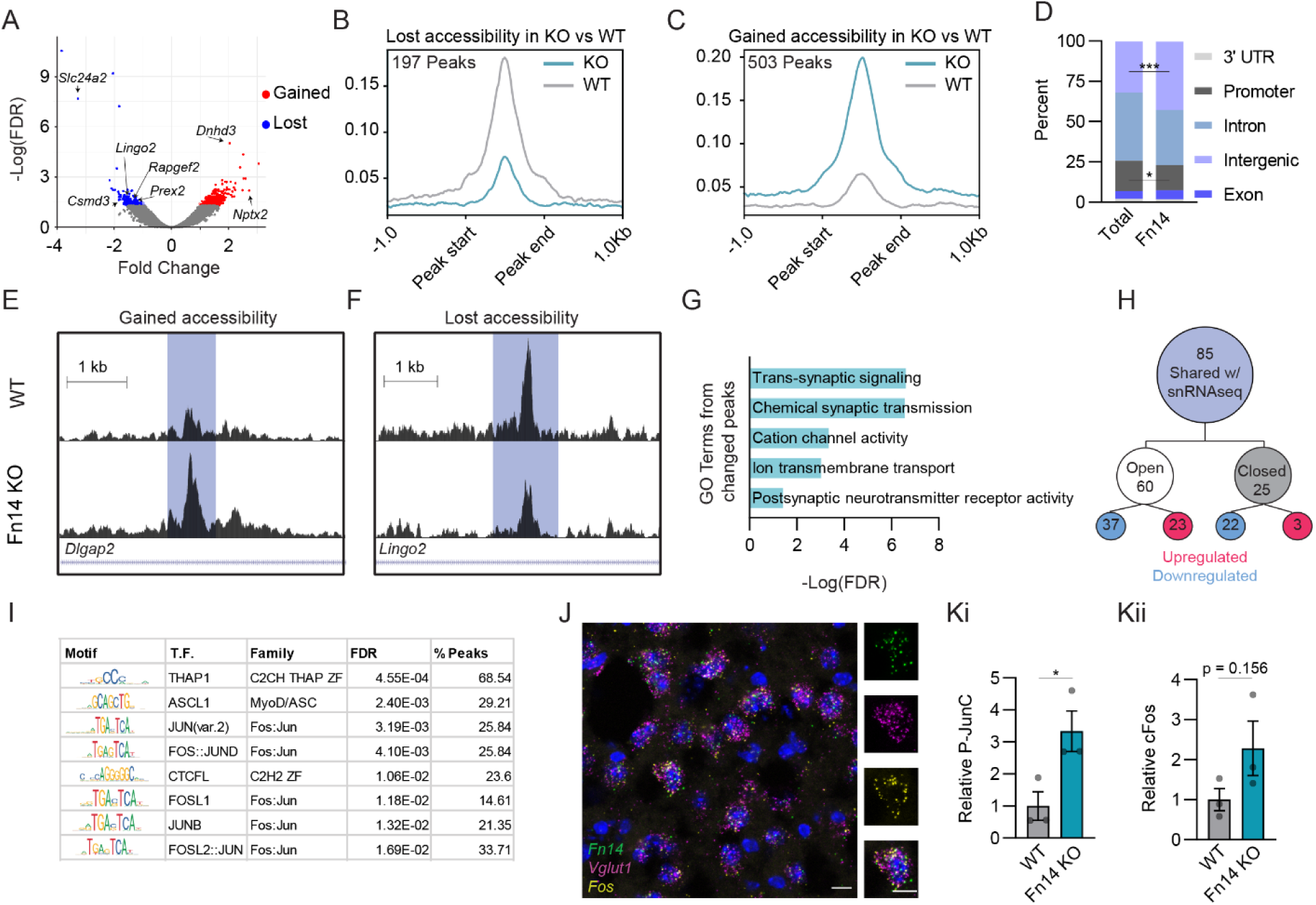
Regulation of chromatin accessibility by Fn14. (A) Volcano plot displaying all ATAC peaks with red points representing genomic regions that are significantly more accessible in Fn14 KO tissue, and blue points representing regions that are significantly less accessible in the KO (FDR< 0.05). (B),(C) Mean read counts per million (CPM; y-axis) across peaks that lost (B) or gained (C) accessibility upon deletion of Fn14. See also figure S5. (D) The location of all peaks (Total) and differentially accessible peaks (Fn14) across the genome. Fischer’s exact test (*p < 0.05, ***p<0.0001). (E) Example of an intronic region within *Dlgap2* that is significantly more accessible in the Fn14 KO compared to WT. x-axis, chromosomal location. Y-axis, average reads. (F) Example of an intronic region within *Lingo2* that is significantly less accessible in the Fn14 KO compared to WT. x-axis, chromosomal location. Y-axis, average reads mapped. (G) GO terms associated with the genes predicted to be regulated by differentially accessible regions. (H) Overlap of differentially accessible peaks and differentially expressed genes (DEGs) in Fn14 KO TC clusters as measured by snRNAseq. (I) Table of predicted transcription factor binding sites within the regions of differentially accessible peaks. % Peaks is defined as the percentage of peaks containing the motif. (J) Fluorescence *in situ* hybridization for *Fn14*, *Vlgut1*, and *Fos* within the dLGN at P28. (Ki),(Kii) Results of protein analysis of AP1 transcription factor activity by ELISA. (Ki), levels of phosphorylated JunC; (Kii) total levels of cFos. Data presented as mean ± S.E.M. with data points representing individual mice (n=3 bioreplicates). Unpaired, 2-tailed student’s T test (*p < 0.05, ***p<0.0001).

In line with dynamic changes in chromatin accessibility occurring largely at non-coding regions, the majority of differentially accessible peaks between Fn14 KO and WT mice were found within intergenic and intronic sequences. We further observed that differentially accessible peaks in the Fn14 KO showed a significant enrichment in intergenic regions compared to all peaks, and a concurrent de-enrichment in promoters (Fig. 3D). In order to assess the nature of the genes that are likely to be impacted by these chromatin changes, we used ChIPSeeker to assign the differentially accessible sites to the genes that they are most likely to regulate (Yu et al., 2015). GO analysis of the genes whose expression is likely to be modified by either chromatin opening or closing downstream of Fn14 revealed a strong enrichment for proteins involved in synaptic transmission, including regulators of postsynaptic neurotransmitter activity, cation channel activity, and trans-synaptic signaling (Fig. 3G). For example, regulatory regions near the genes encoding the neuronal pentraxin Nptx2 which organizes glutamate receptors in the postsynaptic membrane, along with three out of four members of the Dlgap family of excitatory synapse scaffolding proteins, exhibit increased accessibility in the absence of Fn14 (Fig. 3E). Conversely, Lingo2, a cell-surface receptor associated with neurodegeneration, provides an example of a gene associated with a region that is significantly less accessible in the Fn14 KO mouse (Fig. 3F). The observation that sites are both opened and closed in the Fn14 KO is consistent with the up- and downregulation of genes in the absence of Fn14, as well as the ability of both activating and repressing TFs to bind accessible sites.

To determine the extent to which the genes associated with differentially accessible peaks are transcriptionally misregulated in the Fn14 KO mouse, we defined the overlap between the genes associated with these sites and the differentially regulated genes as defined by snRNAseq. We identified 85 genes that are shared between these datasets and might therefore represent the subset of genes whose expression is mediated specifically through the regulation of chromatin downstream of Fn14 (Fig. 3H). Of these 85 sites, 60 were opened in the Fn14 KO mouse and these represented genes whose expression was either up- or down-regulated in the absence of Fn14. The proteins encoded by these shared genes were enriched for effectors of neuronal cytoskeletal remodeling such as Rapgef2, Csmd3, and Prex2, with mutations in Rapgef2 and Csmd3 being strongly associated with familial myoclonic epilepsy (Ishiura et al., 2018). This result is consistent with Fn14 mediating synapse elimination via the structural disassembly of dendritic spines, which could be driven by changes in the expression of these cytoskeletal-regulating genes (Cheadle et al., 2020). Overall, these data highlight that Fn14 shapes neuronal transcription at least in part by establishing the chromatin landscape of developing neurons, and provide a genome-wide resource for uncovering additional Fn14 target genes in the future.

Finally, given that Fn14 is a cell-surface cytokine receptor lacking the ability to directly bind DNA itself, we reasoned that Fn14 likely coordinates changes in transcription and chromatin by engaging intracellular pathways that signal from the cell membrane to the nucleus. If so, then DNA regions that exhibit differential accessibility in the absence of Fn14 may be predicted to bind preferentially to a select cohort of TFs that function downstream of Fn14. We therefore next took advantage of the ATACseq data to identify candidate TFs that may be master regulators of Fn14-dependent transcription by performing a motif enrichment analysis on the DNA sequences underlying peaks that are significantly more or less accessible in Fn14 KO neurons compared to WT (Bailey et al., 2015). This analysis identified a host of TFs predicted to bind genomic regions that are regulated by Fn14, including members of the Specificity Protein/Kruppel-like factor family (e.g. Sp1, Sp3, and KLF5) predicted to bind sites that are more accessible in the Fn14 KO, with Nuclear factor 1 (NFI) TFs predicted to bind sites that are less accessible in the Fn14 KO. Concurrent with NFI motifs being less accessible, *Nfia*, a NFI family member, is downregulated in TC neurons lacking Fn14. Interestingly, focusing specifically on motifs enriched in differentially accessible peaks predicted to regulate the 85 Fn14 gene targets identified in both the snRNAseq and ATACseq datasets, we found that the most enriched motif was predicted to bind Thap1, a TF that is not well understood at the functional level but mutations in which are a major risk factor for the debilitating movement condition dystonia (Houlden et al., 2010). In addition to Thap1, five of the eight significantly enriched motifs are predicted to bind the AP1 family of TFs, including Fos, Jun, and JunB (Fig. 3I). FISH analysis uncovered a strong correlation between *Fn14* and *Fos* expression in the developing and adult dLGN, with the majority (97%) of *Fn14*+ cells being *Fos+* as well, consistent with AP1 factors being involved in the effects of Fn14 on transcription (Fig. 3J). Analysis of AP1 factor activation in Fn14 KO and WT whole brain nuclear extracts by ELISA revealed significant increases in phosphorylated (active) Jun-C and a trending increase in c-Fos mRNA levels in Fn14 KO mice when compared to WT mice (Pulverer et al., 1991)(Fig. 3K). This increase in AP-1 activity alongside the preferential increase in DNA accessibility in the Fn14 KO suggests that Fn14 may function at least in part by repressing AP-1 activity. Altogether, these results characterize Fn14 as a cell-surface receptor that coordinates transcriptional and chromatin changes in neurons during a critical period of SD refinement in the brain. Given that these changes were observed at a time point when Fn14 is required for synaptic refinement, these results further highlight Fn14 as a potential mechanistic link between synapse elimination and epigenomic maturation during postnatal development.

### Microglia mediate chromatin accessibility and synapse elimination during SD refinement

Fn14 is one of many neuronal receptors capable of responding to cytokines expressed by microglia in the healthy brain. The regulation of chromatin accessibility by Fn14 is in line with the possibility that microglia, as a cell class, may engage a plethora of additional immune-related cascades to shape transcription and chromatin organization in developing neurons beyond the TWEAK-Fn14 pathway. To determine whether microglia play a role in organizing chromatin accessibility in developing neurons, we utilized a pharmacological agent, Plexxikon 5622 (PLX), to deplete microglia from the brain selectively during the critical period of SD refinement in the dLGN. PLX depletes microglia by inhibiting the Colony Stimulating Factor 1 receptor which is necessary for microglial survival (Li et al., 2017). Feeding mice chow formulated with PLX beginning at P18 led to the efficient removal of virtually all microglia from the dLGN by P20 and persisted until the end of the experiment between P27 and P32 (Fig. 4A-C; Fig. S5C). While the drug does not provide a high degree of spatial resolution, with depletion occurring not only in the dLGN but across the entire central nervous system, it has the advantage of tight temporal control, allowing us to restrict microglial depletion to the critical period of SD refinement in the dLGN.

**Figure 4.**
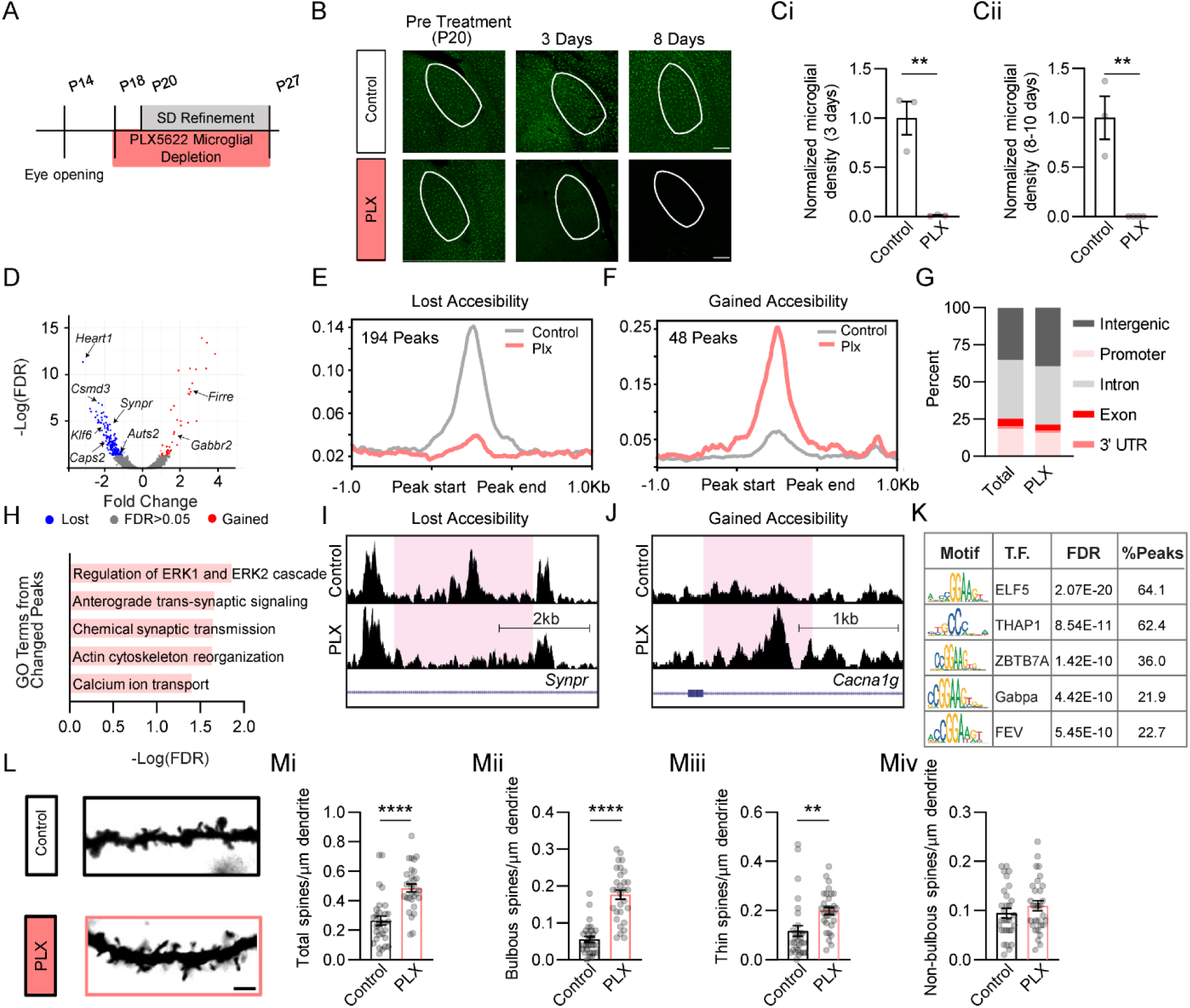
Microglia regulate chromatin accessibility and are required for sensory-dependent refinement in the dLGN. (A) Timeline of retinogeniculate development and the Plexxikon-5622 (PLX) treatment schedule. (B) Confocal images of microglia stained for Iba1 (green) in control and PLX-treated mice before PLX-administration at P20, 3 days into treatment, and 8 days into treatment (white outlines, dLGN. Scale bars, 200 μm). (C) Quantification of the number of microglia in mice fed PLX or control chow confirming near complete depletion of microglia following PLX administration for 3 (Ci) or 8 (Cii) days (n = 3 mice/group). (D) Volcano plot displaying all ATAC peaks with red points representing genomic regions that are significantly more accessible in PLX-treated tissue, and blue points representing regions that are significantly less accessible in PLX-treated tissue (FDR< 0.05). (E),(F) Mean read counts per million (CPM; y-axis) across peaks that lost (E) or gained (F) accessibility upon depletion of microglia. See also figure S5. (G) The location of all peaks (Total) and differentially accessible peaks (PLX) across the genome. (H) GO terms associated with the genes predicted to be misregulated based upon the differentially accessible peaks in PLX-treated mice. (I) Example of decreased accessibility within the intronic region of *Synaptoporin* in microglia-depleted mice. (J) Example of increased accessibility within the intronic region of *Cacna1g* in microglia-depleted mice. (K) Table of the top 5 predicted transcription factor (T.F.) binding sites enriched within differentially accessible peaks. (L) Example images of TC neuron spines from PLX and control mice labeled by Golgi staining. Scale bar, 2 μm. (Mi)-(Miv) Quantifications of the densities of all spines (Mi), bulbous spines (Mii), thin spines (Miii), and non-bulbous spines (Miv) between PLX and control mice. Data presented as mean ± SEM (Control: n =30, PLX: n = 30 spines from 3 mice/condition). Unpaired, 2-tailed student’s T test (**p < 0.01, **** p < 0.0001).

To determine whether microglia are important for organizing chromatin accessibility across the TC neuron genome, we performed ATACseq on the dLGNs of mice at P32 following microglial depletion beginning at P20, alongside age-matched controls fed normal chow in parallel. Of the 192,939 consensus peaks identified in the dataset, 263 were differentially accessible depending upon the presence or absence of microglia (Fig. 4D)(supplemental table 7). To focus on the sites that are most likely to be differentially accessible in TC neurons specifically versus those that were identified due to the removal of microglia from the PLX-treated condition, we filtered out 21 peaks predicted to regulate genes that we found to be microglia-enriched versus neuronally enriched in our Fn14 KO snRNAseq dataset. Analysis of differentially accessible peaks by EnrichR revealed that, after filtering, the sample was enriched for immature neurons (Fig. S5Dii). Overall, this resulted in a final dataset of 194 peaks that were significantly less accessible and 48 peaks that were significantly more accessible in the dLGNs of mice lacking microglia, indicating that microglia tend to increase rather than decrease chromatin accessibility in neurons (Fig. 4E,F; Fig. S5G). The majority of these peaks were found in intronic and intergenic sequences, consistent with these sites representing non-coding DNA elements that regulate neuronal transcription (Fig. 4G).

To define the functions of the genes predicted to be regulated by microglia-dependent changes in chromatin accessibility, we performed GO analysis. We observed a significant enrichment of genes predicted to mediate anterograde trans-synaptic signaling and chemical synaptic transmission (e.g. the presynaptic membrane trafficking regulators *Syntenin-1* and *Caps2,* the synaptic vesicle component *Synaptoporin,* and the ion channel *Cacna1g*) along with several intracellular pathways that convey signals from cell-surface receptors to the nucleus, including the Erk/MapK pathway, which plays an important role in the activation of numerous TFs. Mediators of actin cytoskeleton reorganization (including the autism-susceptibility gene *Auts2* and *Pleckstrin*) were also associated with differentially accessible peaks, consistent with microglia altering the structure of synaptic spines in the dLGN, which we have previously shown to be an important part of SD retinogeniculate refinement (Fig. 3H-J). Motif enrichment analysis of differentially accessible peaks in PLX-treated mice revealed an enrichment of binding sites for Thap1, the TF also predicted to mediate the effects of Fn14 on the genome (Fig. 4K). Nevertheless, comparison of differentially accessible peaks in the PLX versus the Fn14 KO dataset revealed a low degree of overlap with only 12 genes predicted to be regulated by both microglia and Fn14. This observation is consistent with the likelihood that the TWEAK-Fn14 pathway is just one of many signaling cascades engaged by microglia to shape neuronal maturation. Overall, these data highlight that microglial depletion alters chromatin accessibility particularly near genes with critical functions at synapses and in membrane-to-nucleus intracellular signaling.

Although postmitotic neurons undergo epigenomic maturation at the same time as circuits are being actively refined, the functional relationship between these two processes is unknown. We hypothesized that, if the regulation of the epigenome is one method through which microglia refine developing circuits, then microglial depletion should impair not only chromatin accessibility but also the removal of retinogeniculate synapses in response to experience. To test this hypothesis, we quantified retinogeniculate synapses in PLX-treated mice versus mice fed control chow using Golgi-staining and dendritic spine analysis, a read-out that we previously showed to serve as a structural correlate of functional retinogeniculate synapses (Cheadle et al., 2020). While we did not observe overall differences in the length and width of spines between the two conditions, we found that mice lacking microglia maintained 1.8 times more spines overall than mice with microglia (Fig. 4L,Mi). Applying our recent definition of the three major spine populations in the dLGN (Cheadle et al., 2020) revealed that microglia-depleted mice maintained 3.2 times more mature, bulbous-shaped spines and 1.7 times more thin-shaped spines but roughly the same number of non-bulbous spines (Fig. 4Mii-iv). These data indicate that microglia are required for the concurrent patterning of chromatin accessibility and the elimination of synapses in response to sensory experience, suggesting that these two processes may be mechanistically linked.

### Glutamatergic neurons are the predominant expressers of Fn14 throughout the developing and mature brain

TWEAK and Fn14 play a powerful role in SD refinement in the developing dLGN, and the data described above indicate that one mechanism of action is through transcriptional regulation of genes with critical and diverse roles in neurons. Our observations in the dLGN coupled with a recent study describing a role for TWEAK in dampening long-term potentiation in the mature hippocampus (Nagy et al., 2021) led us to explore whether TWEAK-Fn14 signaling may be important for non-visual aspects of brain connectivity and function. To assess this possibility, we first visualized Fn14 expression across the mouse brain using multiplexed FISH (RNAscope) of sagittal sections from mice during peak SD refinement at P28, as well as at P90 after the brain has fully matured. At both P28 and P90, Fn14 expression was restricted to specific neuronal populations across a variety of brain structures, with expression generally increasing along an anterior-to-posterior axis. Fn14 expression was particularly high in the cerebellum where it was largely restricted to the granule cell layer, as well as in the brain stem, thalamus, and select cells in the hippocampus and cortex (Fig. 5A-D). As in the dLGN, the hippocampus exhibited a high degree of co-expression of *Vglut1* and *Fn14* such that ∼80% of *Fn14*+ cells in CA1, CA3, and the dentate gyrus also expressed *Vglut1*. Analysis of *Fn14* mRNA at P90 revealed that *Fn14* expression is retained in glutamatergic neurons across all regions analyzed as the brain matures (Fig. 5E; Fig. S6). As observed in the dLGN at P28, the majority of *Fn14*+ cells in the adult dLGN and hippocampus were also enriched for *Fos* (Fig. 5F,G). Altogether, these results suggest that the roles of TWEAK-Fn14 signaling extend beyond the developing visual system and may be relevant to the mature function of non-visual circuits as well.

**Figure 5.**
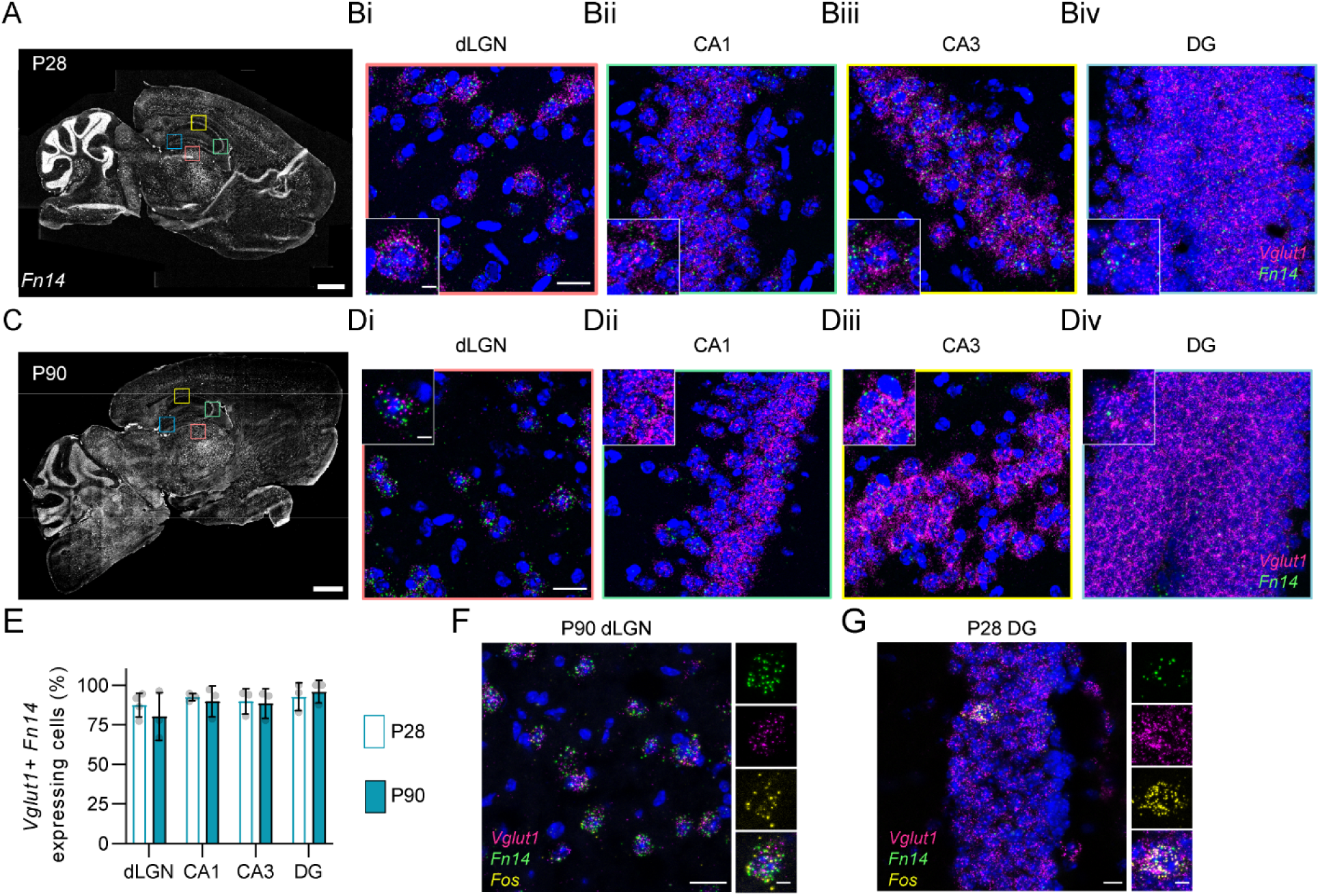
Analysis of Fn14 expression during development and in the adult. (A) Sagittal scan of *Fn14* expression (white) visualized by fluorescence *in situ* hybridization (FISH) in the mouse brain at P28. Boxes correspond with subregions displayed in B. Scale bar, 1 mm. (Bi)-(Biv) High-resolution (63x) confocal images of dLGN (Bi), CA1 (Bii), CA3 (Biii) and dentate gyrus (Biv) at P28. *Fn14* (green), *Vglut1* (magenta), DAPI (blue). Scale bars, 20 μm, and 5 μm for insets. (C) Sagittal scan of Fn14 expression (white) in the brain at P90. (Di)-(Div) High-resolution (63x) confocal images of dLGN (Di), CA1 (Dii), CA3 (Diii) and dentate gyrus (Div) at P28. *Fn14* (green), *Vglut1* (magenta), DAPI (blue). Scale bars, 20 μm, and 5 μm for insets. (E) Quantification of the percentage of *Fn14*+ cells that represent *Vglut1*+ excitatory neurons. 89.3% of *Fn14*+ cells are also *Vglut1*+ across brain regions and timepoints (datapoints representing averaged values from individual mice, n = 3) . See also figure S6. (F),(G) High-resolution (63x) confocal images of *Fn14*, *Vglut1*, and *Fos* RNA at P90 in dLGN (F) and at P28 in the DG (G).

### Fn14 is dispensable for learning but required for memory task proficiency

While the roles of microglial cytokine signaling in the healthy brain have been most extensively studied in the context of synapse elimination during development, several recent studies have suggested that these pathways also play a role in maintaining circuit homeostasis, excitatory/inhibitory balance, and even complex behaviors in the adult (Badimon et al., 2020; Nguyen et al., 2020; Wang et al., 2020). We speculate that some of the mechanisms underlying synapse refinement during development may also underlie synaptic plasticity in the mature brain, particularly in the case of microglia and cytokine molecules like Fn14 given their critical roles in chromatin accessibility. In light of the expression of Fn14 in excitatory neurons of the hippocampus, a brain region that is critical for both learning and memory and the function of which can be assayed robustly using well-defined behavioral tests, we next sought to determine whether Fn14 facilitates hippocampal function in adult mice.

We tested the roles of Fn14 in learning and memory using two well-established tasks: cued fear conditioning (CFC) and Morris water maze (MWM). In the CFC task, we examined the abilities of Fn14 KO and WT littermates to associate both an auditory cue and a spatial context with a paired aversive foot shock (Fig. 6A). During the initial conditioning phase, when the foot shock was accompanied by a tone (75 dB; 2000 Hz) and a novel arena (striped walls and floor grating), both Fn14 KO and WT mice exhibited a stereotyped freezing response reflecting fear of the shock. Similarly, when mice of both genotypes were placed into a novel context (a round arena with polka dotted walls) without a tone, they exhibited low levels of freezing. However, loss of Fn14 resulted in a trending decrease in freezing when re-exposed to the paired context but not the tone (context (-) tone condition), and a significant decrease in freezing when the novel context was paired with the associated tone (novel context +tone)(Fig. 6B). These results suggest that Fn14 KO mice did not sufficiently retain the memory that the tone was paired with a foot shock.

**Figure 6.**
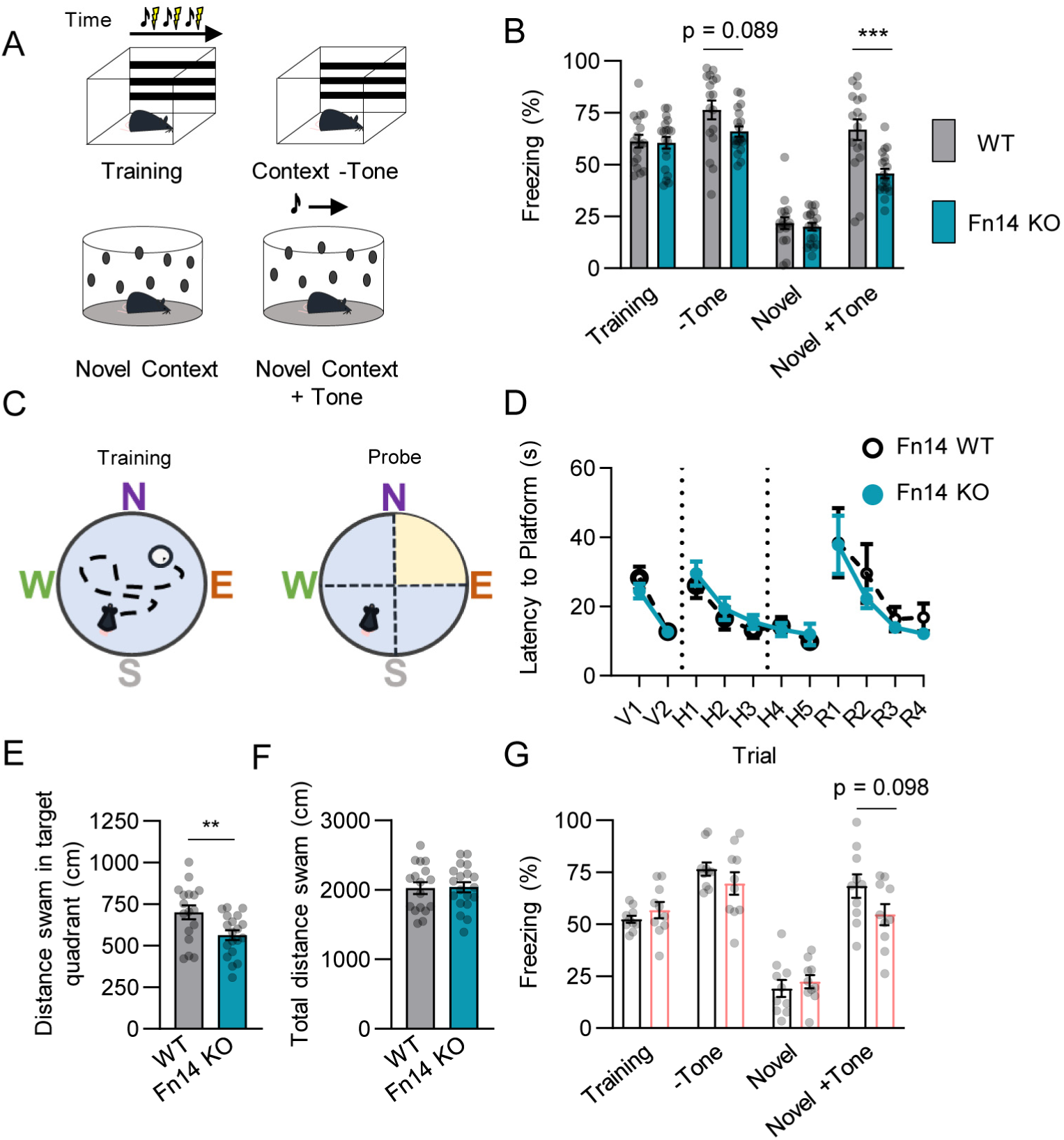
Loss of Fn14 impairs memory but not learning in multiple behavioral tasks. (A) Diagram of the cued fear conditioning (CFC) paradigm. (B) Quantifications of percentage of time spent freezing across all conditions (repeated measures ANOVA, trial: p < 0.0001; genotype: p < 0.05; subject, trial x genotype: p < 0.0001). Bonferroni corrected multiple comparisons WT versus KO for Context (-) tone: p = 0.089; Novel context + tone: p < 0.001. (C) Diagram of Morris Water Maze (MWM) training and probe trials. (D) Latency to goal platform swam during MWM trials (repeated measures ANOVA with Šídák’s multiple comparisons test: Visible; genotype: p > 0.05, trial: p < 0.0001, trial x genotype: p > 0.05, Hidden; genotype: p > 0.05, trial: p < 0.0001, trial x genotype: p > 0.05, Reverse; genotype: p > 0.05, trial: p < 0.001, trial x genotype: p > 0.05). (E) Distance swam by mice in the target quadrant during probe trials. (F) Total distance swam by mice during probe trail. See also figure S7. (G) Percentage of time spent freezing for control-chow-administered (n = 8; gray bars) and PLX-administered (n = 8; pink bars) mice. Repeated measures ANOVA with Bonferroni corrected multiple corrections: trial: p < 0.0001, treatment p > 0.05, trial x treatment p > 0.05. Data presented as mean ± S.E.M. with data points representing individual mice. (E) and (F) Analyzed with student’s T test (**p < 0.01, ***p < 0.001).

Because impairments in the CFC task could reflect functional changes in the amygdala or the frontal cortex in addition to the hippocampus, to determine whether these deficits may reflect hippocampal dysfunction specifically, we next assessed the effect of loss of Fn14 on a more purely hippocampal-dependent spatial learning task, the MWM. In this task, the mice were placed in a round pool with each cardinal direction being marked by a distinctive shape to allow for spatial mapping of the arena (Fig. 6C). During the initial visible training stage, WT and Fn14 KO mice were both able to effectively locate a visible goal platform. After mice were trained to perform the task, the water in the pool was made opaque, and the goal platform submerged, forcing mice to orient themselves using the spatial cues to locate the goal platform rather than the platform itself. In all trials in which the platform was hidden, WT and Fn14 KO mice learned to find the platform equally well as revealed by their similar latencies to reach the platform and the lengths of the paths that they took to reach it (Fig. 6D; Fig. S7A,B). Thus, as also demonstrated by the results of the CFC task, loss of Fn14 does not have a strong observable effect on learning.

To specifically assess memory function, we next tested whether, after a period of 24 hours, the mice remembered the location of the hidden platform. When the platform was removed from the pool in probe trials, WT mice swam a significantly greater distance in the quadrant where the platform was previously hidden than Fn14 KO mice, suggesting that WT mice were able to remember the spatial location of the platform while mice lacking Fn14 were unable to remember the goal location (Fig. 6E). This deficit was not caused by a motor impairment, as WT and Fn14 KO mice swam an equal distance overall during the probe trial (Fig. 6F). Following the probe trials, the goal platform was reintroduced into the pool, but now in the opposite quadrant of the arena. Just as in the hidden trials, both WT and Fn14 KO mice were able to locate and learn the new reversed goal zone equally well, again suggesting that Fn14 does not affect learning. Rather, the specific deficits observed in the probe trials suggest that loss of Fn14 impairs long-term memory in this hippocampal-dependent task.

In order to determine whether microglia are also important for these tasks, we next tested PLX-administered, microglia-depleted mice in both CFC and MWM. As with Fn14 KO mice, PLX mice froze to the same extent as control mice during initial conditioning to the averse stimulus as well as when placed in the novel context. PLX-administered mice also froze to an equal extent as control mice when the mice were re-exposed to the conditioned context but not the tone. Interestingly, there was a trending decrease in freezing in PLX mice when they were re-exposed to the conditioned tone in a novel context (p = 0.0989), which was the condition with the greatest effect size in Fn14 KO mice (Fig. 6G). This result suggests that this aspect of the memory deficits in mice lacking Fn14 may be related to aberrant signaling from microglia. Nevertheless, microglial depletion did not significantly affect performance in the MWM task, confirming that some aspects of Fn14 function likely do not require microglia (Fig. S7D-I). Overall, these data suggest that Fn14 signaling contributes to memory but not learning, consistent with our previous findings that synapses in the Fn14 KO mouse fail to properly strengthen (Cheadle et al., 2018).

### Loss of Fn14 exacerbates PTZ-induced seizures

In combination with recent studies demonstrating a requirement of TWEAK and Fn14 for visual circuit development and hippocampal plasticity, our findings that Fn14 mediates the expression of neuronal transcription and chromatin accessibility and is required for memory suggest that this pathway is critical for mature brain function. Thus, we next sought to determine whether Fn14 regulates overall (i.e. gross) neural activity in adult mice *in vivo*. Using electroencephalogram (EEG) probes implanted into the dorsal skull, we quantified the effect of loss of Fn14 on brain activity over a 48-hour period (Fig. 7A). To ensure the mice had similar baseline physiology, the animal’s overall locomotor activity and body temperature were monitored. Fn14 KO and WT mice exhibited equal body temperature and activity levels over the recording session (Fig. S8), similarly to what we observed in the MWM task (Fig. S6). Correspondingly, we found that there was no difference in average EEG spectral power between Fn14 KO and WT mice in delta (1-4 Hz), theta (4-8 Hz), alpha (8-12 Hz), beta (12-30 Hz), low gamma (30-60 Hz) and high gamma (60-90 Hz) frequency bands (Fig. 7B). Therefore, as implicated by the normal performance of mice lacking Fn14 in the learning phases of both the CFC and the MWM, loss of Fn14 does not affect baseline neural activity on a gross level.

**Figure 7.**
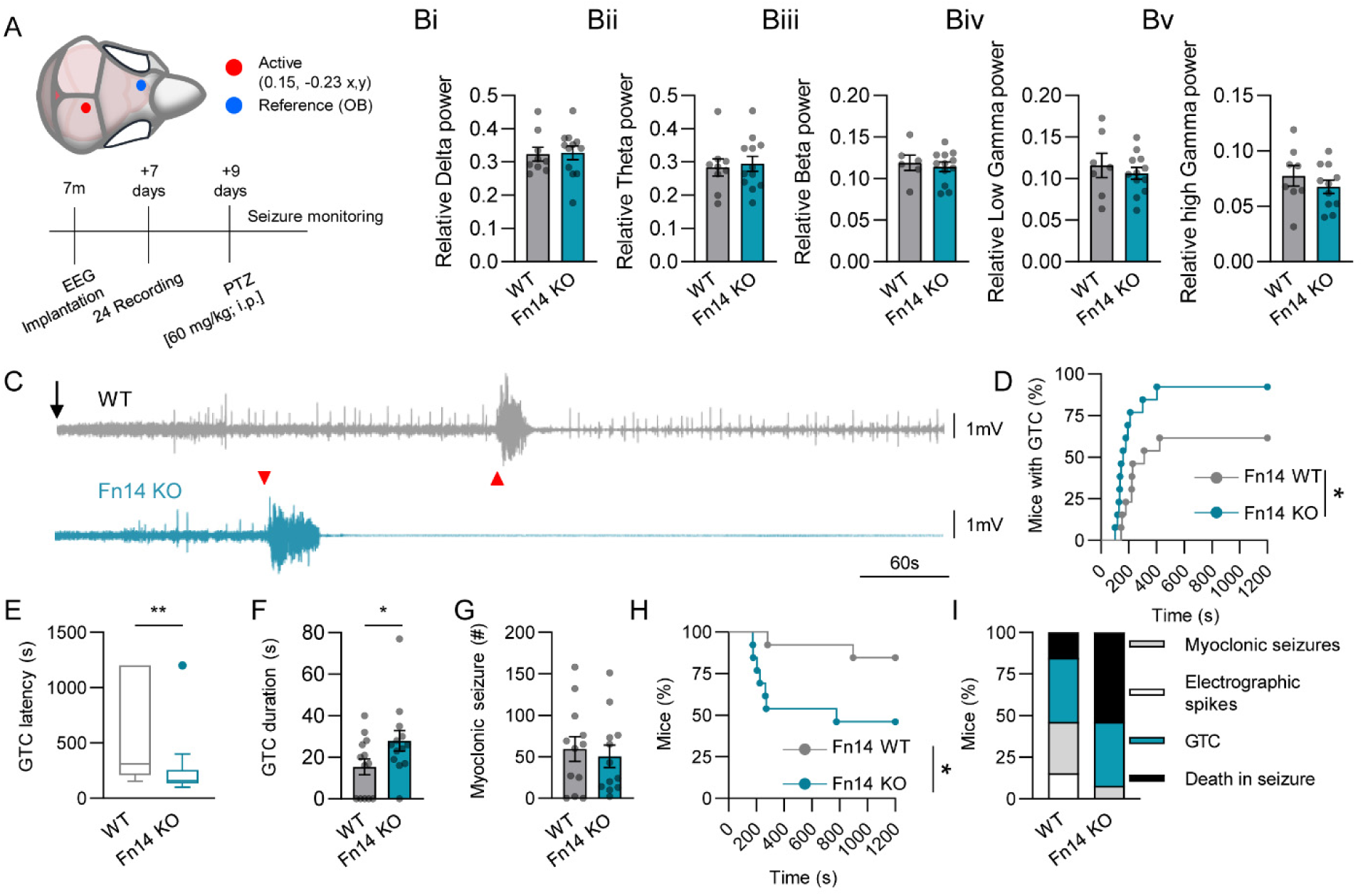
Loss of Fn14 confers seizure susceptibility and increased mortality following exposure to PTZ. (A) Schematic of electroencephalogram (EEG) electrode placement and the experimental timeline. (Bi)-(Bv) Averaged EEG spectral power across frequency bands were equal between Fn14 knockout (KO) and WT mice (WT n = 11, Fn14 KO n = 12, unpaired Student’s t-test p > 0.05). See also figure S8. (C) Example EEG traces from WT (gray) and Fn14 KO (teal) mice after PTZ injection (black arrow). Red triangles indicate the onset of general tonic clonic (GTC) seizures (WT: latency = 311 s, duration = 19.8 s; Fn14 KO: latency = 159 s, duration = 35 s). The Fn14 KO mouse died shortly after the GTC, demonstrated by the elimination of signal following the seizure. (D) Percentage of mice that had GTC seizures relative to the time course of the experiment (WT; n = 13 median = 311 s, Fn14 KO; n = 13, median = 159 s; Log-Rank test; *p < 0.05). (E) Latency between PTZ injection and GTC onset (Mann-Whitney test; **p < 0.01). (F) Duration of GTCs (unpaired Student’s T-test; *p < 0.05). (G) Number of PTZ-induced myoclonic seizures (Mann-Whitney, p > 0.05). (H) Mortality rate of Fn14 KO and WT mice following PTZ administration. Log-Rank test; *p < 0.05. (I) The fraction of mice presenting with electrophysiological spikes (white), myoclonic seizures (grey), GTCs (teal), or death as their worst PTZ-induced outcome. Data presented as mean ± S.E.M. with data points representing individual mice or as the percentage of subjects where applicable.

Consistent with Fn14 functioning at least in part downstream of microglial cytokines, studies using PLX to deplete microglia also found that PLX-treated mice did not exhibit baseline changes in EEG spectral power across the range of relevant bands overall. Rather, microglial depletion resulted in increased seizure risk and exacerbated seizure outcomes when mice were challenged with a neural stimulant (Badimon et al., 2020). Given that mice without microglia maintain an excess of synapses due to deficits in synapse elimination which are also observed in Fn14 KO mice (Cheadle et al., 2018), we wondered whether loss of Fn14 would lead to a similar susceptibility to seizures. To determine whether loss of Fn14 confers seizure risk, we analyzed the responses of Fn14 KO mice to the GABA_a_ antagonist pentylenetetrazole (PTZ), a convulsant agent used to elicit seizures (Van Erum et al., 2019). Intraperitoneal injection of PTZ (60 mg/kg) into Fn14 KO and WT mice time-locked with EEG recordings demonstrated profound differences in the responses of mice of each genotype to PTZ. Upon PTZ injection, Fn14 KO mice were more likely than WT littermates to develop general tonic clonic (GTC) seizures, and Fn14 KO mice developed GTCs at a shorter latency than WT mice (Fig. 7C-E). Furthermore, GTC seizures had a longer duration in Fn14 KO mice (Fig. 7F) when compared to GTCs in WT mice. Concurrently with the increase in GTC severity, Fn14 KO mice had a significantly higher mortality rate after PTZ challenge than WT mice (Fig. 7H). Interestingly, Fn14 KO mice experienced significantly fewer myoclonic seizures than WT mice (Fig. 7G), potentially due to the higher mortality of Fn14 KO mice. Lastly, loss of Fn14 led to a worse overall seizure phenotype as scored by a combination of their recorded behavior, EEG activity, and mortality, suggesting that loss of Fn14 confers an increased susceptibility to acutely induced seizures, similarly to published data on microglia depleted mice (Fig. 7I). Altogether, these functional data reveal that, although Fn14 KO mice do not exhibit overt deficits in brain activity at baseline, upon increasing excitatory tone through PTZ-mediated disinhibition, loss of Fn14 exacerbates seizure severity and worsens seizure outcomes. Taken together, these findings highlight new roles for microglia and Fn14 in mediating neuronal transcription and chromatin accessibility, leading to the refinement of circuits during development and the function of circuits in the mature brain.

## Discussion

Over the past several years, it has become increasingly evident that cytokine signaling pathways play essential roles in the organization of neural circuits in the developing and mature brain (Ferro et al., 2021). However, we are still in the early stages of defining the downstream mechanisms through which these pathways shape neuronal connectivity in the long-term, particularly during SD phases of synaptic refinement. Here, we derive significant insights into how a TNF-family cytokine signaling axis that mediates inflammation outside of the brain, the TWEAK-Fn14 pathway, also controls a postnatal stage of circuit refinement through the coordination of gene expression and chromatin remodeling in excitatory TC neurons of the dLGN. We additionally show that microglia simultaneously regulate patterns of chromatin accessibility and synapse elimination during the critical period of SD retinogeniculate refinement, corroborating that these aspects of brain development result in part from microglia-neuron interactions. Furthermore, we demonstrate that expression of Fn14 extends to excitatory neurons in multiple non-visual structures in the developing and mature brain, suggesting that Fn14 coordinates multiple aspects of brain function, many of which likely remain to be identified. Supporting this possibility, behavioral analysis revealed that Fn14 is dispensable for learning but that loss of Fn14 impairs memory. This observation is consistent with previous reports of Fn14’s effect on synaptic plasticity during development (Cheadle et al., 2018), and in the adult hippocampus (Nagy et al., 2021). Finally, we demonstrate that, while mostly normal at baseline, seizure activity in mice lacking Fn14 is excessively heightened upon exposure to a convulsant, leading to a high mortality rate in mice lacking Fn14. Overall, this multi-disciplinary study elucidates the roles of Fn14 signaling in the brain from the regulation of genes encoding effectors of synaptic and epigenomic remodeling during development to brain-wide activity and cognitive function in the adult.

Our data support a model (Fig. 8) in which membrane-bound Fn14, either in response to or independently from microglial signals including TWEAK, induces intracellular signaling pathway(s) to mediate gene expression in the neuronal nucleus. While the nature of these signaling cascades and the TFs that they activate remain unclear, our observation that chromatin accessibility increases at AP-1 TF binding sites in the Fn14 KO and that AP-1 factors show heightened activation in the KO suggest that Fn14 may function at least in part by repressing AP-1 activity. In any case, we propose that pathways downstream of Fn14 upregulate the transcription of genes encoding synaptic organizers which, upon translation, localize at synapses to promote their maturation and/or disassembly. In addition to this cohort of synapse-regulating genes, a second cohort of Fn14 targets encodes molecules that modify histones and chromatin in the neuronal nucleus, altering the accessibility of *cis*-regulatory regions of the genome. These alterations in chromatin patterning lead to the expression of an additional cohort of genes, the products of which function predominantly to induce structural synaptic remodeling and plasticity. Given that TWEAK and Fn14 KO mice also display increased expression of some transcripts, it is likely that TWEAK and Fn14 can mediate synaptic refinement through the negative regulation of synaptic remodelers as well. We speculate that the Fn14-regulated synapse-organizing proteins that *are not* associated with chromatin changes promote the refinement of developing synapses that takes place during the critical period between P20 and P30. We further speculate that the synaptic proteins that *are* associated with changes in chromatin remain poised to facilitate changes in synapse number and function into maturity. In other words, the synaptic proteins that are not associated with chromatin changes may regulate synapses transiently while those that are associated with chromatin changes—which are likely to be longer-lasting— remain poised to remodel synapses in the long-term.

**Figure 8.**
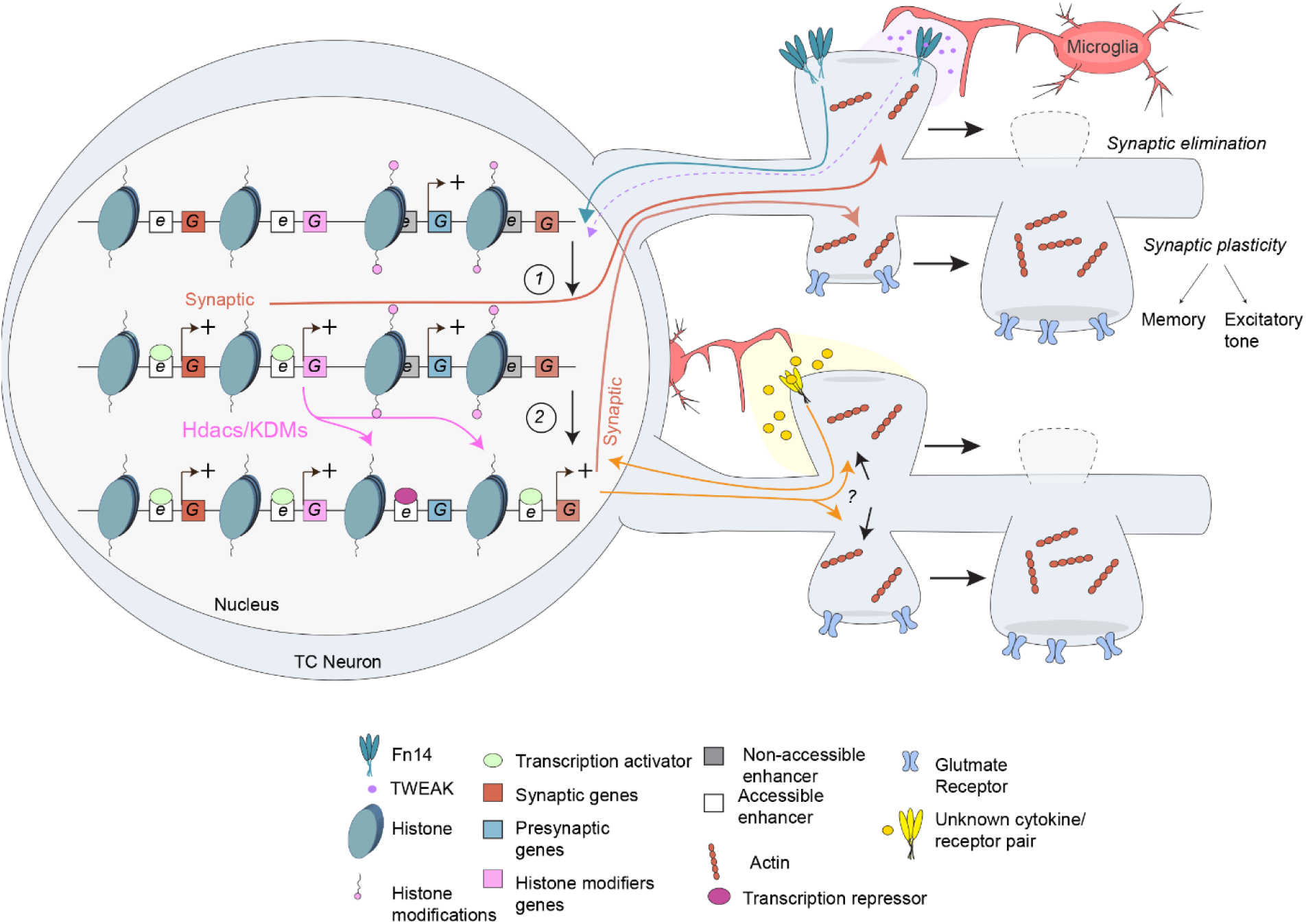
Proposed model of Fn14 and microglial regulation of transcription and chromatin accessibility in neurons. Signaling downstream of Fn14 at the cell membrane, either in response to TWEAK binding or through self-association, activates intracellular cascades that control transcription in the neuronal nucleus. Fn14 and TWEAK upregulate the expression of genes encoding mediators of two essential processes: (a) synaptic function, and (b) chromatin remodeling (*1*). We speculate that the synaptic regulators induced by Fn14 (orange) localize to synapses to directly facilitate their elimination, while the chromatin and histone modifiers, i.e. histone deacetylases and demethylases (Hdacs and KDMs, pink), remain in the nucleus to reshape chromatin accessibility near an additional cohort of genes that encode mediators of synaptic function (*2*). The synaptic regulators whose expression corresponds with Fn14-dependent changes in chromatin accessibility may represent factors that remain poised to remodel synapses across the lifespan. Microglia also control chromatin accessibility in neurons near genes encoding mediators of synapse development and function via mechanisms that have yet to be defined, the majority of which are likely to be distinct from TWEAK-Fn14 signaling. We speculate that the misregulation of transcription and chromatin accessibility in mice lacking Fn14 contributes to the decreased memory function and seizure susceptibility observed in Fn14 KO mice.

Consistent with this model, critical periods of postnatal brain development correspond with a transient developmental process of epigenomic maturation that involves permanent changes to the genomic architecture, impairments in which can lead to neurodevelopmental disorders such as intellectual disability and autism (Frank et al., 2015; Stroud et al., 2017; Stroud et al., 2020). While much remains to be uncovered about the mechanisms of Fn14 function, our model begins to explain why memory formation, a process that requires new gene transcription as well as acute structural and functional changes at mature synapses (Alberini and Kandel, 2014), is disrupted in the absence of Fn14. At the same time, given that the homeostatic downscaling of synapse number and strength are associated with heightened levels of neural excitation and that these adaptations can protect the brain from excitotoxicity (Lignani et al., 2020), the lasting misregulation of synaptic plasticity genes capable of adapting to increased firing may also begin to explain the increased seizure susceptibility of mice lacking Fn14. These results, along with our finding that microglia shape chromatin organization and synaptic refinement during a SD period of circuit development, present a new framework for how cytokine signaling pathways regulate neural circuit connectivity through mechanisms beyond the local remodeling of individual synapses.

While establishing the control of transcription and chromatin as a new method through which Fn14 regulates brain development and function, this work raises several questions about TWEAK-Fn14 signaling in the brain and microglia-neuron communication as a whole. First, given the limited overlap between the genes that are misregulated in the absence of Fn14 versus TWEAK, the extent to which TWEAK contributes to Fn14-dependent gene regulatory mechanisms remains incompletely understood. Because constitutive genetic loss of TWEAK across the lifespan may induce compensatory mechanisms making it difficult to assess the targets of TWEAK *in vivo*, future experiments overexpressing TWEAK in the brains of Fn14 KO and WT mice according to a more acute paradigm may provide additional insights into how TWEAK mediates neuronal transcription through binding to Fn14. Second, little is known about the downstream intracellular signaling partner(s) engaged by Fn14 to control neuronal transcription. While NfκB was a prime candidate based upon the activation of this pathway by Fn14 in the periphery, our ATACseq data did not reveal an enrichment of NfκB binding sites among the *cis*-regulatory regions that are controlled by Fn14. On the contrary, our data suggest that Fn14 function in the brain may instead involve transcriptional regulators such as THAP1 and AP1 family members. Another outstanding question is whether the chromatin changes that are elicited by Fn14 and microglia during development are transient or last into maturity, as our model speculates the latter. In addition, given that microglia induce chromatin changes that are largely distinct from those elicited by Fn14, it is likely that additional cytokine signaling pathways linking microglia to neuronal chromatin organization remain to be discovered. Addressing these open questions in the future could lead to important insights not only into the roles of cytokine signaling and microglia in brain development, but also into how impairments in the epigenomic maturation of neurons may give rise to neurological diseases emerging across the lifespan.

Compared to the roles of microglia and cytokines in brain development, less is known about the ongoing functions of cytokines in the mature brain. This is a critical gap in knowledge to address given that neuroinflammation is associated with a broad range of neurological disorders emerging after development, such as multiple sclerosis, stroke, and neurodegeneration. In addition to the role of TWEAK and Fn14 in establishing neural circuits, data also suggest that TWEAK-Fn14 signaling may be relevant in neurodegenerative disease, particularly in the context of Alzheimer’s disease (AD), which is thought to be exacerbated by the aberrant re-activation of refinement mechanisms in the mature brain (Hammond et al., 2019; Hong et al., 2016). Fn14 expression is heightened in brain tissue from individuals with AD, and a recent study showed that inhibiting TWEAK-Fn14 signaling using a TWEAK-blocking antibody rescues synaptic plasticity deficits in a well-established mouse model of AD (Nagy et al., 2021). Our data support a potential role for heightened TWEAK-Fn14 signaling in disorders of cognition and memory, including AD, by demonstrating that Fn14 is expressed in the hippocampi of adult mice, and that loss of Fn14 results in impaired memory in multiple hippocampal-dependent tasks. In conclusion, this work provides evidence that Fn14 specifically, and microglia more broadly, impact neural transcription and chromatin accessibility, and that the dysregulation of these developmental pathways likely contributes to impaired behavioral outcomes in adult mice.

## Methods

### Animal models

All experiments were performed in compliance with protocols approved by the Institutional Animal Care and Use Committee (IACUC) at Harvard Medical School and Cold Spring Harbor Laboratory. The following mouse lines were used in the study: C57Bl/6J (the Jackson Laboratory; Bay Harbor, MA; JAX:000664); B6.Tnfrsf12a^tm1(KO)Biogen^ (Fn14 KO)(Jakubowski et al., 2005); and B6.Tnfsf12^tm1(KO)Biogen^ (TWEAK KO)(Dohi et al., 2009). TWEAK and Fn14 KO mice were generously provided by Dr. Linda Burkly at Biogen (Cambridge, MA) and are subject to a Material Transfer Agreement with Cold Spring Harbor Laboratory. Analyses were performed on equal numbers of male and female mice at <u>p</u>ostnatal days (P)27 or P90. No sex differences were observed in the study.

### Plexxikon 5622 administration

Chow formulated to contain 1200 mg/kg free base Plexxikon 5622 (PLX; Chemgood, Inc.; Glen Allen, VA) was fed to mice between P18 and P27. Control mice were fed chow containing the same ingredients and produced in parallel with PLX-formulated chow, but not containing PLX. All chow was produced and irradiated by Research Diets, Inc. (New Brunswick, NJ) and stored at 4°C. PLX treatment did not cause observable changes in animal health or behavior. As per our standard protocol, mice were fed on the chow *ad libitum* and the investigator provided all husbandry for the mice during treatment.

### Golgi staining

Golgi staining was performed with the FD Rapid GolgiStain kit (FD Neurotechnologies, Inc., Columbia, MD) according to the manufacturer’s protocol. Following sample processing, dendritic segments were traced in x, y, and z planes using a Zeiss (White Plains, NY) Axioskop microscope (63X objective) and Neurolucida (Microbrightfield Bioscience; Hong Kong, China). Spines were categorized based upon parameters determined in a previous study (Cheadle et al., 2020). Example images of spines were obtained on an Olympus BX63 fluorescence microscope with a 100X objective.

### Immunofluorescence

To validate microglial depletion, PLX-fed and control mice were perfused with ice cold PBS (Gibco; Dublin, Ireland) and 4% paraformaldehyde (PFA), then the whole brains were harvested and post-fixed for 12 hours. After fixation, tissue was incubated in 15% and then 30% sucrose solutions before being embedded in OCT (-80°C). Embedded tissue was sectioned coronally at 25 μm thickness onto Superfrost Plus slides using a Leica (Danvers, MA) CM3050 S cryostat. Sections were then washed in PBS and blocked in blocking solution (PBS adjusted to 5% normal goat serum [NGS] and 0.3% Triton X-100 [TX-100]) for 1 hour at room temperature before being incubated in primary antibody solution containing Rabbit-anti-Iba1 (Wako 019-19741, [1:1500]; Richmond, VA) diluted in PBS with 5% NGS and 0.1% TX-100 (probing solution). Sections were incubated in primary antibody overnight at 4°C. The next day, sections were washed 3 times for 10 minutes per wash in PBS before incubation in secondary antibody Alexafluor 488 Goat-anti-rabbit (Thermo Fisher 150077; [1:500]; Waltham, MA) diluted in probing solution for 2 hours at room temperature. Sections were then washed in PBS, covered with DAPI fluoromount-G (SouthernBiotech; Birmingham, AL), and cover-slipped.

Confocal images (20X) were acquired using a LSM 710 Zeiss microscope and the number of microglia/dLGN was quantified using ImageJ. A total of 3 mice per condition and a minimum of two images per mouse were analyzed. In some cases, we increased the contrast and subtracted the background across an entire image to clarify the features of interest that are presented in the figures. Other than contrast enhancement applied across the entire image, no other manipulations to images were made.

### Single-nucleus RNA-sequencing

#### Tissue preparation, nuclear capture, and next-generation sequencing

Fn14 KO and WT littermate mice and TWEAK KO and WT littermate mice at P27 were euthanized and their brains harvested in ice cold PBS (Gibco). Coronal sections of 300 μm were made on a VT1000S vibratome (Leica) in ice cold PBS and the dLGNs were micro-dissected using a Nikon SMZ10A brightfield dissection microscope. Dorsal-LGN tissue from three mice per condition per replicate was pooled and homogenized in HB buffer containing .25 M sucrose and (in mM): 25 KCl, 5 MgCl_2_, 20 Tricine-KOH, pH 7.8, and 2.5% Igepal-630 (Sigma). HB was adjusted to contain 1 mM DTT, .15 mM spermine, and .5 mM spermidine along with phosphatase and protease inhibitors (Roche). All steps were performed on ice. Following homogenization, the homogenate was layered atop a 30%-40% iodixanol gradient and centrifuged in an SW-41 swing bucket rotor (Beckman) at 10,000*g* for 18 minutes. Small volumes of <u>B</u>ovine <u>S</u>erum <u>A</u>lbumin (BSA; Sigma) and RNAsin (Promega, Madison, WI) were included in HB and iodixanol solutions. Nuclei were recovered from the 30%-40% iodixanol interface and individual nuclei were captured within microfluidic droplets alongside barcoded hydrogels via the inDrops approach (Zilionis et al., 2017). After cell encapsulation, primers were released by UV exposure. Libraries of 2500 – 3000 nuclei were generated for each sample (one library per bioreplicate for two bioreplicates total of the Fn14 WT/KO line (four libraries total) and two libraries per bioreplicate for four bioreplicates of the TWEAK WT/KO line (16 libraries total). Indexed libraries for each mouse line were independently pooled and sequenced twice on a Next-Seq 500 (Illumina; San Diego, CA) with sequencing parameters set at Read 1, 54 cycles; Read 2, 21 cycles; Index 1, 8 cycles; Index 2, 8 cycles.

#### Data processing and analysis

Reads were mapped against a custom transcriptome built from Ensemble GRCm38 genome and GRCm38.84 annotation using Bowtie 1.1.1, after filtering the annotation gtf file (gencode.v17.annotation.gtf filtered for feature_type = “gene”, gene_type = “protein_coding” and gene_status = “KNOWN”). Read quality control and mapping were performed. Sequence reads were linked to individual captured molecules by unique molecular identifiers (UMIs). Default parameters were used unless stated explicitly. These steps were previously described (Cheadle et al., 2018; Macosko et al., 2015).

#### Quality control, cell clustering, and differential gene expression analysis

All Fn14 KO/WT nuclei were combined into a single dataset and the same was done for TWEAK KO/WT nuclei, because these experiments were performed separately. The Fn14 and TWEAK datasets were analyzed in parallel using the same approach. First, the R package DoubletFinder (https://github.com/chris-mcginnis-ucsf/DoubletFinder)(McGinnis et al., 2019) was applied to each sample separately to predict and remove potential doublets from the dataset. This step is crucial for ensuring that none of the nuclei in the final dataset actually represent two nuclei encapsulated within the same droplet, which could skew the analysis. This step removed roughly 6% of the nuclei from the dataset. Next, Seurat version 3 was used to further filter out nuclei that may represent surviving doublets or that may be dead or dying cells by removing all nuclei from the dataset that did not contain between 250 and 3,000 UMIs and also removing cells containing greater than 5% mitochondrial read counts (Stuart et al., 2019). In Seurat v3, the data were log normalized and scaled to 10,000 transcripts per cell, and variable genes were identified using default parameters. We limited the analysis to the top 30 principal components (PCs). Clustering resolution was set between 0.5 and 1.0. Clusters containing fewer than 100 cells were removed from the dataset, as were clusters expressing more than one known marker gene. Known marker genes were used to assign clusters to cell types as follows: excitatory thalamocortical neurons, *Slc17a6* and *Prkcd*; all inhibitory neurons, *Gad2*; dLGN-surrounding inhibitory neurons, *Penk*; dLGN-resident inhibitory neurons, *Chrnb3*; oligodendrocytes, *Olig1*; oligodendrocyte precursor cells, *Pdgfra*; astrocytes, *Aqp4* and *Aldoc*; microglia, *P2ry12*; endothelial cells, *Cldn5*; and pericytes, *Vtn*. Excitatory relay neurons, the primary focus of the study, represented ∼51% of 11,586 total nuclei in the Fn14 WT/KO dataset and 47% of 50,004 nuclei in the TWEAK WT/KO dataset.

To identify genes that were differentially expressed between KO and WT conditions within a given cell class, we used Monocle 2 (https://github.com/cole-trapnell-lab/monocle2-rge-paper), an R package specialized for quantitative analysis of single-cell data (Qiu et al., 2017). Genes whose differential gene expression false discovery rate (FDR) was less than 0.05 (FDR < 0.05) were considered statistically significant. General descriptions of differentially expressed genes are also based upon a fold-change cutoff of 1.25-fold.

Gene ontology (GO) analyses were performed based upon PANTHER term enrichment on genes that were significantly downregulated or upregulated in TWEAK or Fn14 KO nuclei (or the genes whose expression was predicted to be misregulated in the ATACseq data) with FDR < 0.05 regardless of absolute fold change (Mi et al., 2013).

### ATAC-sequencing and analysis

#### Tissue isolation and next-generation sequencing

Fn14 KO and WT littermates (P27) were euthanized, and their brains harvested and flash frozen (1 mouse per replicate, 1 male and 1 female per condition). At the time of processing, the brains were thawed and a coronal slice containing both dLGN and hippocampus was dissected. To study the effects of microglial depletion on chromatin accessibility, age-matched PLX-chow and control-chow fed mice were euthanized (1 male and 1 female mouse per replicate, 2 replicates per condition) and their dLGNs micro-dissected. PLX tissue then entered processing directly. Both the Fn14 and PLX experiments followed the same preparation after dissection. Samples remained on ice throughout, unless otherwise noted. The tissue was homogenized in 1.4 mL ice cold ATAC buffer (AB) containing (in mM): 10 Tris-HCl pH 7.4, 10 NaCl, 3 MgCl_2_, .5 Spermidine, .15 Spermine, 1 DTT, 0.1% Igepal, 1X Phosphatase inhibitor cocktails 2 and 3 (Sigma P5726, P0044), and 1X Protease inhibitor tablets (Sigma-Aldrich; St. Louis, MO). Homogenization was performed by douncing with a tight pestle 40 times. The samples were then spun at 500 rcf for 10 minutes. The pellet was resuspended in 1.4 mL fresh AB and dounced again 12 times. Next, the nuclei were washed by spinning for 5 minutes and resuspending in fresh AB. 50,000 nuclei were aliquoted according to the nuclear concentration obtained using the Countess II FL Automated Cell Counter (Thermo Fisher). Aliquots were spun once more at 500 rcf, and then resuspended in 20 µL of DNAse free water. Illumina Tagment DNA kit (cat. 20034197) enzyme and buffer were added to the DNA samples, which were then incubated at 37°C for 30 minutes. The DNA was then purified using the Qiagen MinElute PCR Purification kit (Germantown, MD). Libraries were prepared for each sample with standard protocols and the NEBNext High-Fidelity PCR Master Mix (Ipswich, MA) for 11 amplification cycles, and then purified once again. Next, at room temperature, the samples were diluted 1:1 with 50% glycerol, ran along a 2% agarose electrophoresis gel, and stained with ethidium bromide in order to excise bands between 100 and 1000 bp. The excised DNA was purified using the Qiagen MinElute Gel Extraction kit based upon the manufacturer’s instructions. Libraries within an experiment (Fn14 KO/WT or PLX/control chow) were pooled and subjected to paired end sequencing on an Illumina NextSeq with a High Output Kit.

#### Data processing and differential peak analysis

Both Fn14 and PLX ATAC-sequencing data were analyzed in the same manner. The ENCODE ATAC-seq pipeline was run applying default parameters. With the pipeline’s output, BAM and MACS2 narrowPeaks files were fed into DiffBind (Ross-Innes et al., 2012) to generate a dba object from which a consensus peak set was created by recentering ± 250 bp around the highest reads (dba.count, summits = 250). Next, a contrast was automatically generated (dba.contrast, categories = DBA_TREATMENT) and differentially accessibly peaks were called using edgeR.

#### Annotation, motif analysis, and plots

The differentially accessible peaks generated with DiffBind were annotated using the ChIPSeeker package (annotatePeak, default parameters) and the UCSC mm10 known genes. For the PLX samples, we filtered out the genes that were twice as enriched in WT microglial snRNAseq clusters than in WT thalamocortical snRNAseq clusters. This filtered set of peaks was used for subsequent analysis and plots. Transcription factor motif enrichment was performed using MEME-AME (McLeay and Bailey, 2010). Differentially accessible peak sequences were inputted into the MEME-AME (https://meme-suite.org/; McLeay et al., 2010) program using the JASPAR core non-redundant vertebrates database with background control data consisting of randomly shuffled peak sequences derived from all (differentially accessible and non-differentially accessible) peak sequences within a given dataset. To identify cell types that may be enriched within our differentially accessible peaks, the EnrichR program (Chen et al., 2013) using the PanglaoDB Augmented 2021 database was used. DeepTools (Ramirez et al., 2014) was used to generate bigwig tracks (binsize = 1, normalize = CPM), heatmaps and peak profiles. Volcano plots were generated using the R package ggplot2. Two peaks in the Fn14 volcano plot (annotated as Tnfrsf12a: Log_2_(FC) = -4.97, -Log(FDR) = 27.01; and Ddr1: Log_2_(FC) = -6.44, -Log(FDR) = 21.13) were removed to better represent the majority of the data.

### Quantification of AP-1 binding to DNA

P27 Fn14 KO and WT whole brain tissue (n = 3 mice/genotype) was collected and flash frozen in liquid nitrogen. Tissue was later thawed and homogenized in RIPA buffer (VWR; Atlanta GA) via agitation on ice for 30 minutes before centrifugation at 23,000 x *g* for 10 minutes. 5 microliters of the insoluble fraction were then diluted in Complete Lysis Buffer (Active Motif; Carlsbad, CA) and nuclear protein concentration was determined using a Bradford assay (Bio-rad; Hercules, CA). Once nuclear proteins were diluted to equal concentrations in Complete Lysis Buffer, 20μg of sample was then used to quantify binding of c-Fos and phosphorylated Jun-C (P-Jun-C) to oligonucleotides consensus binding sites for AP-1 family members according to the manufacturer’s instructions. Briefly, nuclear extracts were added to a pre-coated 96-well plate, and antibodies against P-Jun-C and c-Fos were added and the plate was incubated for 1 hour at room temperature. After washing each well, an HRP-conjugated secondary antibody against P-Jun-C or c-Fos was added and the plate was incubated at room temperature for another hour. After washing off the unbound secondary antibody, each colorimetric reaction was developed and subsequently stopped using Stop solution. Absorbance at 450 nm was measured for protein binding within 5 minutes of addition of Stop solution with 650 nm as a reference. Technical replicates (n = 2/sample) were averaged and data was normalized to WT samples.

### RNA isolation and qPCR

Fn14 KO and WT littermate mice and TWEAK KO and WT littermate mice at P27 were euthanized and their brains were bisected and flash frozen using liquid nitrogen in 1 mL of Trizol (Ambion; Naugatuck, CT) and kept at -80°C until processing. Tissue was then homogenized using a motorized tissue homogenizer (Thermo Fisher) in a clean, RNAase-free environment. Once homogenized, 200 μL of chloroform was added to each sample and, after thorough mixing, samples were centrifuged at 21,000x*g* for 15 minutes for phase separation. The colorless phase was then collected and combined with equal volume of 70% ethanol and used as input in the RNeasy Micro kit (Qiagen), where we followed the manufacturer’s instructions. RNA concentration was then determined using a nanodrop (ND 1000; NanoDrop Technologies inc.; Wilmington, DE), and once RNA samples were diluted to equal concentrations, samples were converted into cDNA using SuperScript™ III First-Strand Synthesis System (Thermo Fisher) following the manufacturer’s instructions. Specific genes were then amplified (forward and reverse primers can be found in supplemental table 5) and detected using Power Up Sybr Green (Thermo Fisher) in a Quant Studio 3 Real-Time PCR system (Thermo Fisher). Crossing threshold (Ct) values were calculated using the QuantStudio program and relative expression, 2^-ΔΔCt^, was calculated using *GAPDH* as a reference control.

### Single-molecule fluorescence in situ hybridization (FISH)

Sagittal sections of 20 – 25 μm thickness were made using a Leica CM3050 cryostat, collected on Superfrost Plus slides, and stored at -80°C. Multiplexed single-molecule FISH was performed using the RNAscope platform (Advanced Cell Diagnostics [ACD]; Newark, CA) according to the manufacturer’s protocol for fresh-frozen sections (multiplexed detection kit version 1). Commercial probes obtained from ACD detected the following genes: *Tnfrsf12a* (*Fn14*), *Slc17a7* (*Vglut1*), *Gad1*, *Penk*, and *Chrnb3*.

For quantification of *Fn14*, *Slc17a7*, and *Gad1* transcripts/cell, 60X confocal images were acquired using a LSM 710 Zeiss microscope. A total of 3 mice per condition and a minimum of two images per mouse were analyzed. *Fn14* expression was quantified using an ImageJ macro built in-house (code: www.cheadlelab.com/tools). Briefly, the DAPI channel was thresholded and binarized, and subsequently expanded using the dilate function. This expanded DAPI mask was then passed through a watershed filter to ensure that cells that were proximal to each other were separated. This DAPI mask was then used to create cell specific ROIs, where each ROI was considered a single cell. Using these cell-masked ROIs, the number of FISH puncta were counted using the 3D image counter function within imageJ, within a given ROI. ROIs were classified with the following criteria: ROIs containing 3 or more *Fn14* molecules were considered positive for *Fn14*, and for markers (*GAD1* and *Vglut1*) cells were considered positive if there were 5 or more marker molecules present within a given ROI.

### Behavior

#### Cued Fear Conditioning

On training day, subjects were placed into a square fear-conditioning arena of 24(w)x20(d)x30(h) cm equipped with a shock grid floor and acrylic walls patterned with horizontal black and white bars 2 cm in width. Subjects were allowed to acclimate to the arena for 4 minutes before data acquisition. During training, mice were presented with three 20 second tones (75 dB; 2000 Hz) followed by a 2 second foot shock (0.5 mA) with variable inter-trial intervals totaling 5 minutes. After training, subjects were returned to their home cages for 24 hours before being tested in familiar and novel contexts. For familiar context (the paired context without the cued tone; Context (-) tone) subjects were re-acclimated to the test arena for 5 minutes without receiving tone cues or shocks to reduce freezing to non-tone cues. After testing freezing in the Context (-) tone condition and on the same day, subjects were exposed to a novel context (circular arena 30(w) x 30(h) cm, with clear acrylic floor and polka-dot walls) for 3 minutes to habituate the mice to the novel context before freezing examined. Mice were then returned to their home cages for 24 hours before being re-exposed to the novel context, but were then re-presented with the cued tone (75 dB; 2000 Hz) for three minutes during acquisition. Freezing was calculated using Ethovision XT v. 15 (Nodulus; Wageningen, Netherlands) where freezing was measured using activity detection set to 300 ms and data was presented as freezing over the trial time.

#### Morris Water Maze

Each training trial consisted of four 90 s sub-trials in which each subject’s starting position was pseudo-randomized to each of the four cardinal directions in a 137 cm wide water bath containing 24°C clear water filled up to 25 cm from the rim of the tub. The cardinal directions were marked on the wall of the tub with 20 cm diameter symbols. Subjects were initially trained over two trials, where the goal platform was raised 0.5 cm above the water line and was marked with a bright flag for increased visibility (visible trial). Each trial ended either after the trial time expired, or after the subject correctly found and stayed on the goal platform for more than 5 seconds. If a mouse did not find the platform within 90 seconds, it was gently moved to the platform and left there for 5 seconds. The day following visible platform training, the goal platform was submerged (0.5 cm below the water line) and moved to a different quadrant. Subjects were tested on the hidden platform over 5 consecutive trials spanning 48 hours. On the fourth day (probe trial) the goal platform was removed from the testing arena and subjects were placed facing the wall opposite of the previous goal platform’s position. Subjects were allowed to swim for a total of 60 s before being removed from the arena. On reversal trials (4 trials), the goal platform remained submerged, but was moved to the opposite end of the arena. Subjects started the reverse trials facing the furthest wall and were allowed to search for the goal platform for 90 s. If the subject failed to find the goal platform, the subject was oriented in the correct direction and guided to the goal platform before being removed from the arena. Latency to goal platform, distance swam, and subject position was collected using Ethovision XT v. 15 (Nodulus).

### EEG recordings and PTZ seizure induction

#### EEG telemetry unit implantation

Mice were implanted with wireless telemetry units (PhysioTel ETA-F10; Data Sciences International [DSI]; Holliston, MA) under sterile techniques per laboratory protocol as previously described. Under anesthesia, a transmitter was placed intraperitoneally, and electrodes were threaded subcutaneously to the cranium. After skull exposure, haemostasis, and identification of the cranial sutures bregma and lambda, two 1-mm diameter burr holes were drilled over the right olfactory bulb (reference) and left occipital cortex (active). The epidural electrodes of the telemetry units, connected to the leads of the transmitter, were placed into the burr holes, and secured using stainless steel skull screws. Once in place, the skull screws were covered with dental cement. Mice were subcutaneously injected 0 and 24 hours post-operatively with 5 mg/kg meloxicam for analgesia. After 1 week of recovery, mice were individually housed in their home cages in a 12-h light–12 h dark, temperature and humidity-controlled chamber with *ad libitum* access to food and water.

#### Baseline and PTZ seizure induction

After a 24-h acclimation period, one-channel EEG was recorded differentially between the reference (right olfactory bulb) and active (left occipital lobe) electrodes using the Ponemah acquisition platform (DSI). EEG, core-body temperature, and locomotor activity signals were continuously sampled from all mice for 48 h along with time-registered videos. At the end of baseline acquisition, all mice were provoked with a convulsive dose (60 mg/kg; i.p.) of the GABA_a_ receptor antagonist pentylenetetrazole (PTZ; Sigma-Aldrich, Co.) to measure seizure susceptibility and evaluate seizure thresholds (Dhamne et al., 2017; Yuskaitis et al., 2018; Zullo et al., 2019). Mice were continuously monitored for clinical and electrographic seizure activity for 20 minutes.

#### Data analysis

All data were processed and analyzed using Neuroscore software (DSI). Baseline EEG was analyzed for spontaneous seizure activity, circadian biometrics, and spectral power band analysis (Dhamne et al., 2017; Yuskaitis et al., 2018). Relative spectral power in delta (1-4 Hz), theta (4-8 Hz), alpha (8-12 Hz), beta (12-30 Hz), low gamma (30-60 Hz) and high gamma (60-90 Hz) frequency bands of the baseline EEG were calculated using the fast Fourier transform (FFT) technique.

PTZ-induced seizure activity was broadly scored on a modified Raccine’s scale as only electrographic spikes (score: 1), myoclonic seizures (score: 3), generalized tonic-clonic seizures (GTC; score: 5) and death (score: 6). Per mouse, number of myoclonic seizures, latency and incidence of GTC seizures, number of GTCs, and total duration of GTC were recorded. Mice without seizures were assigned a time of 20 min at the end of the PTZ challenge observation period.

### Blinding

Experimenters were blinded to conditions at all stages of analysis. For immunofluorescence, FISH, Golgi staining, and snRNAseq, one experimenter harvested the tissue and assigned it a randomized label before providing the blinded tissue to another experimenter for analysis. After data acquisition and processing, the data were plotted in Graphpad (San Diego, CA) by L.C. or A.F. after which the samples were unblinded. Similar approaches were used for the behavioral and EEG experiments.

### Reagents and Resources

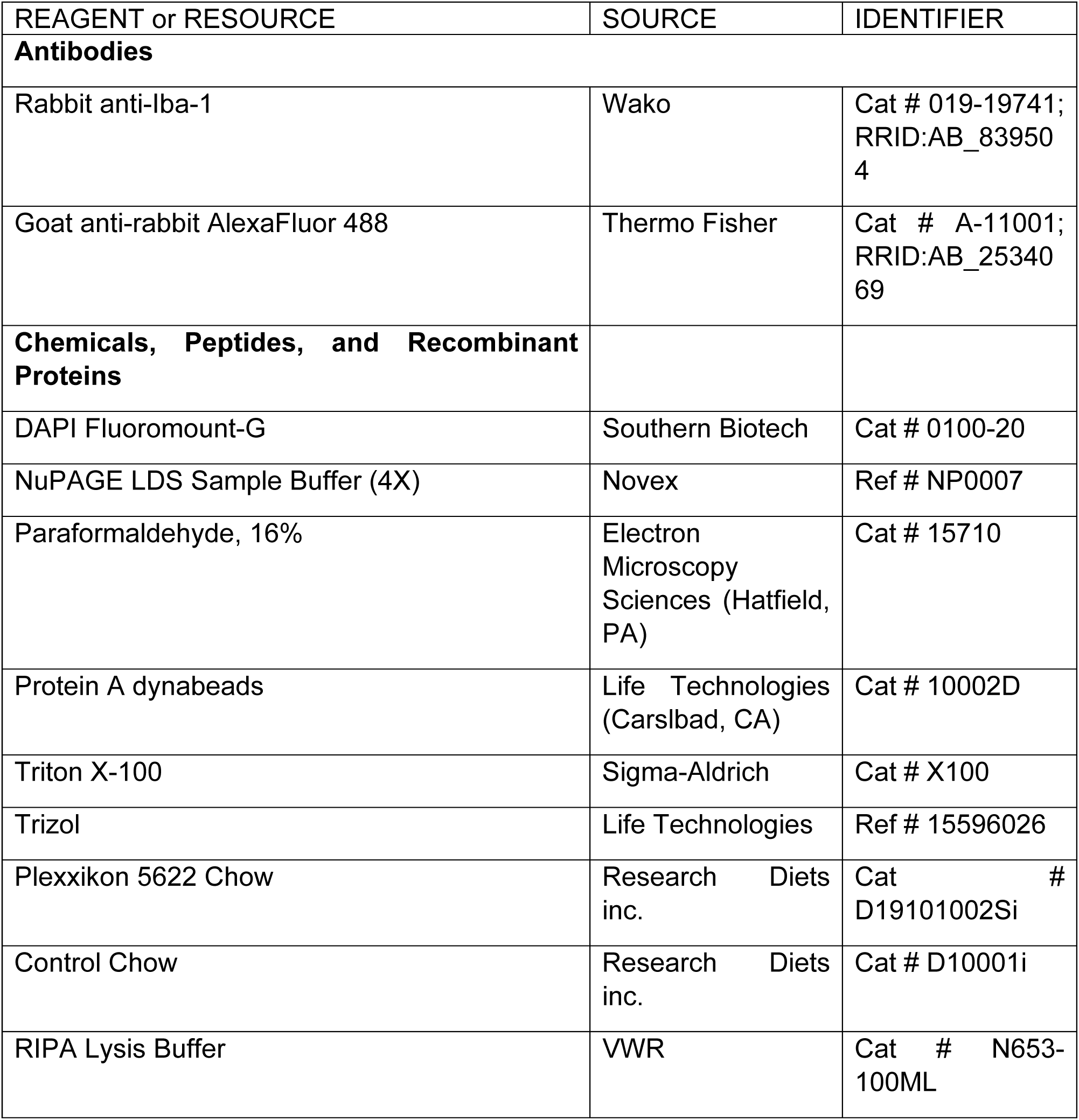
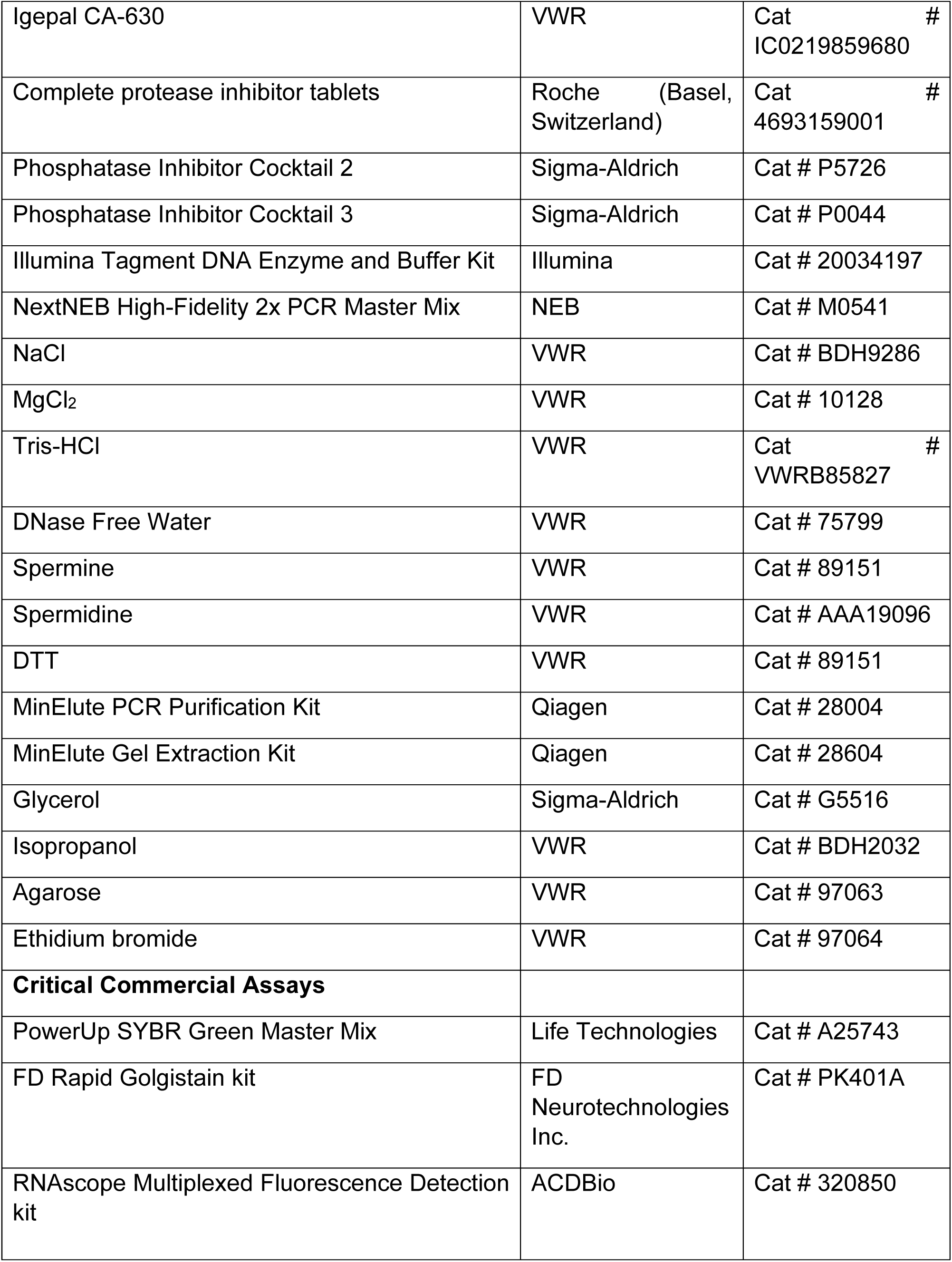
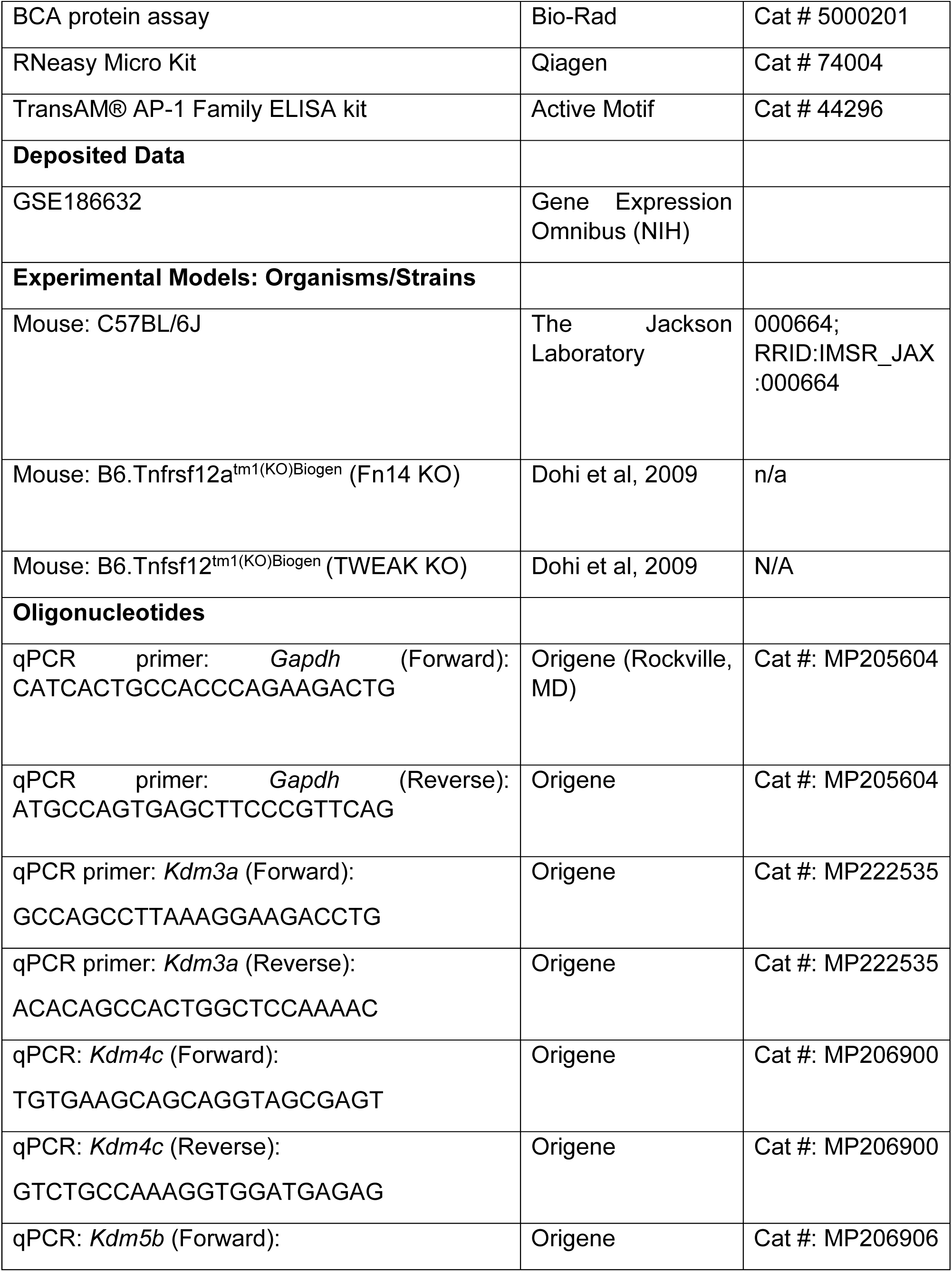
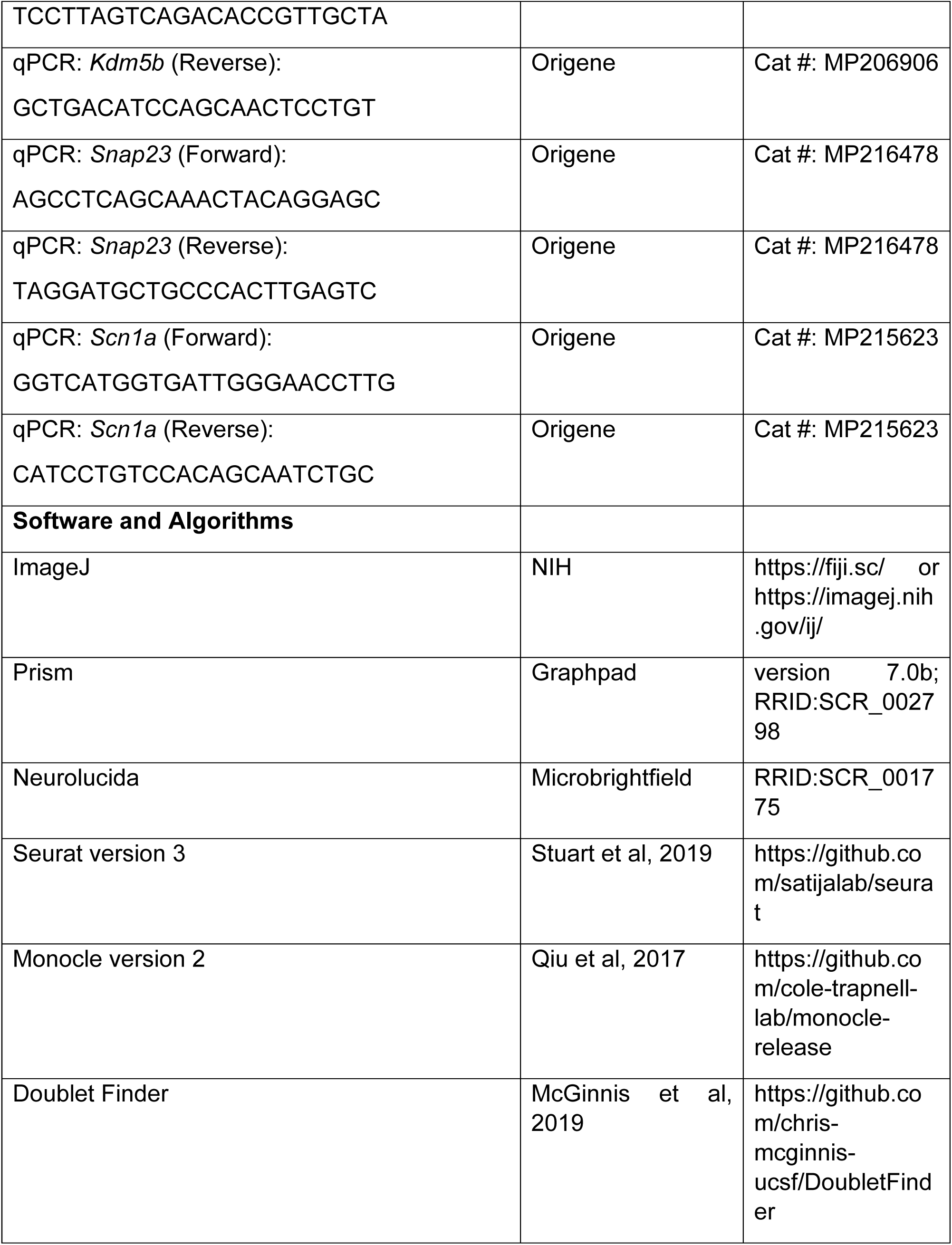
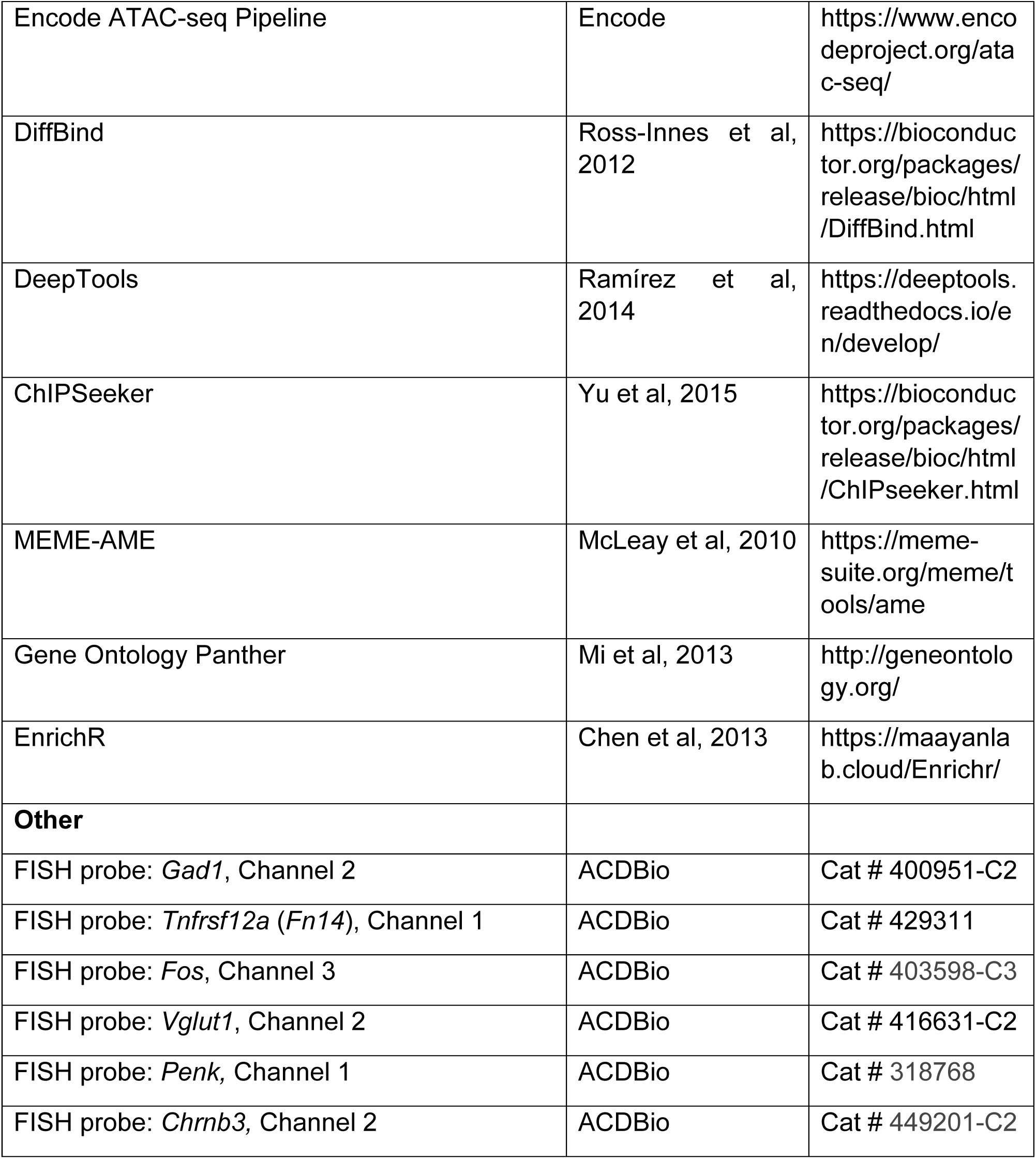

## Acknowledgements

We acknowledge Drs. Linda C. Burkly (Biogen), Lisa D. Boxer (Harvard Medical School), Marty Yang (Harvard Medical School), Sameer Dhamne (Boston Children’s Hospital), Jessica Tollkuhn (Cold Spring Harbor Laboratory), Timothy Cherry (Seattle Children’s Hospital), Jacqueline Barker (Drexel University), Elizabeth Pollina (Harvard Medical School), Marija Cvetanovic (University of Minnesota) and Cody Walters (Nature Communications) for thoughtful discussions and feedback on the study. Work in this study was aided by the Single-Cell core facility at Harvard Medical School and the Behavior and Physiology core at Boston Children’s Hospital. This work was supported by a R00 MH120051-4 (NIMH), a McKnight Scholar Award, a Rita Allen Scholar Award, and a Klingenstein-Simons Fellowship in Neuroscience (to L.C.).

## Author Contributions

L.C. conceptualized the study. L.C., A.F., U.V., and Y.S.S.A. performed experiments. L.C. analyzed snRNAseq and spine data. A.F. analyzed FISH, behavioral, and EEG data. U.V. analyzed qPCR and immunostaining under the guidance of A.F. Y.S.S.A. performed ATACseq analyses, motif enrichment analyses, and tissue processing. L.C. and A.F. wrote the manuscript with input from U.V. and Y.S.S.A.

