## Supplemental Figures 1 - 8 for "Microglia and Fn14 regulate transcription and chromatin accessibility in developing neurons"

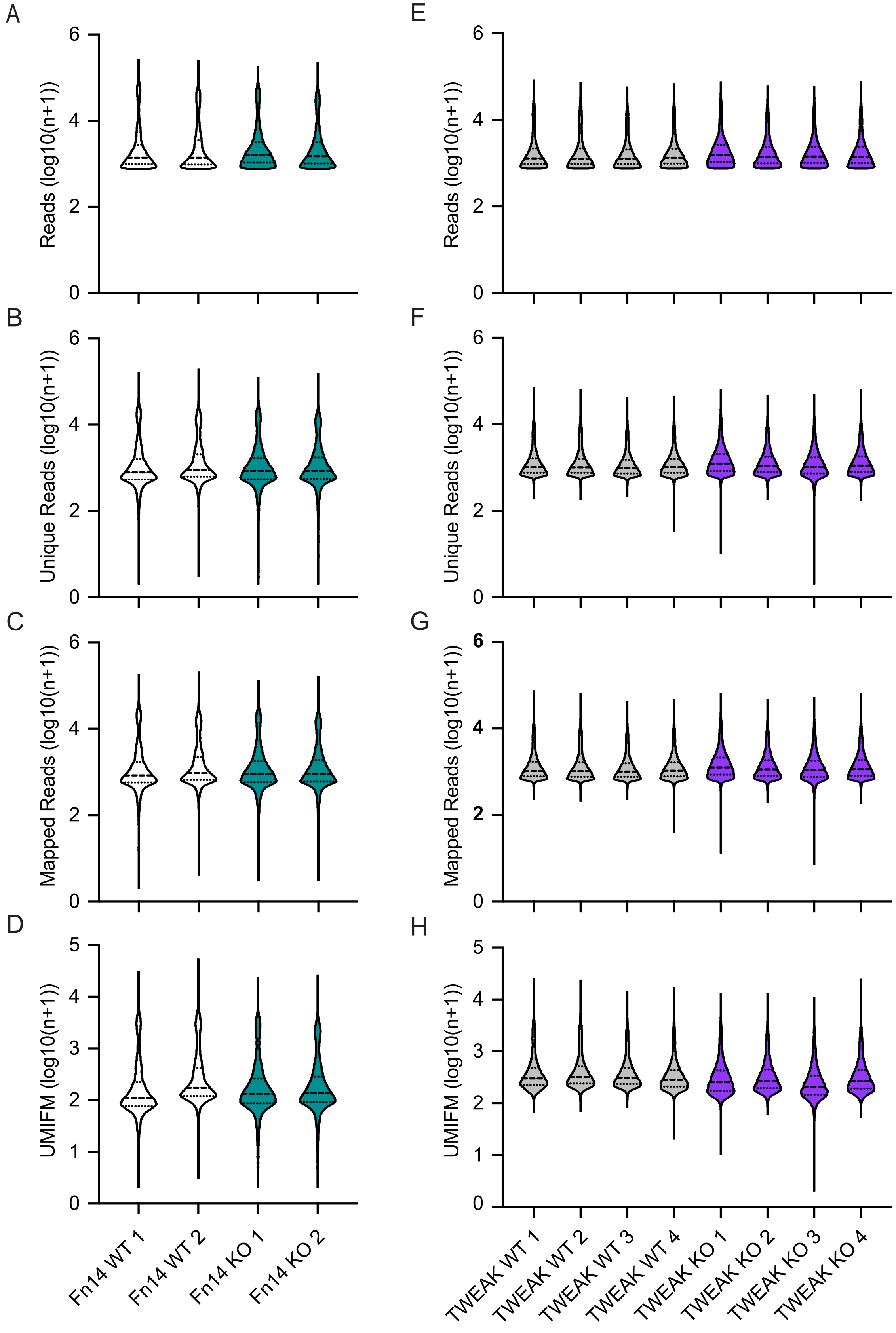


**Figure S1 (Related to figure 1). Quantitative sequencing metrics by sample prior to filtering**. Values expressed as log10(n + 1) where n equals normalized transcripts per cell. For Fn14 KO dataset: (A) Total reads per cell; (B) Unique reads per cell; (C) Mapped reads per cell; (D) Unique Molecular Identifier Filtered Mapped reads (UMIFM) per cell. For TWEAK KO dataset: (E) Total reads per cell; (F) Unique reads per cell; (G) Mapped reads per cell; (H) UMIFM per cell. Dashed lines, median. Dotted lines, quartiles.

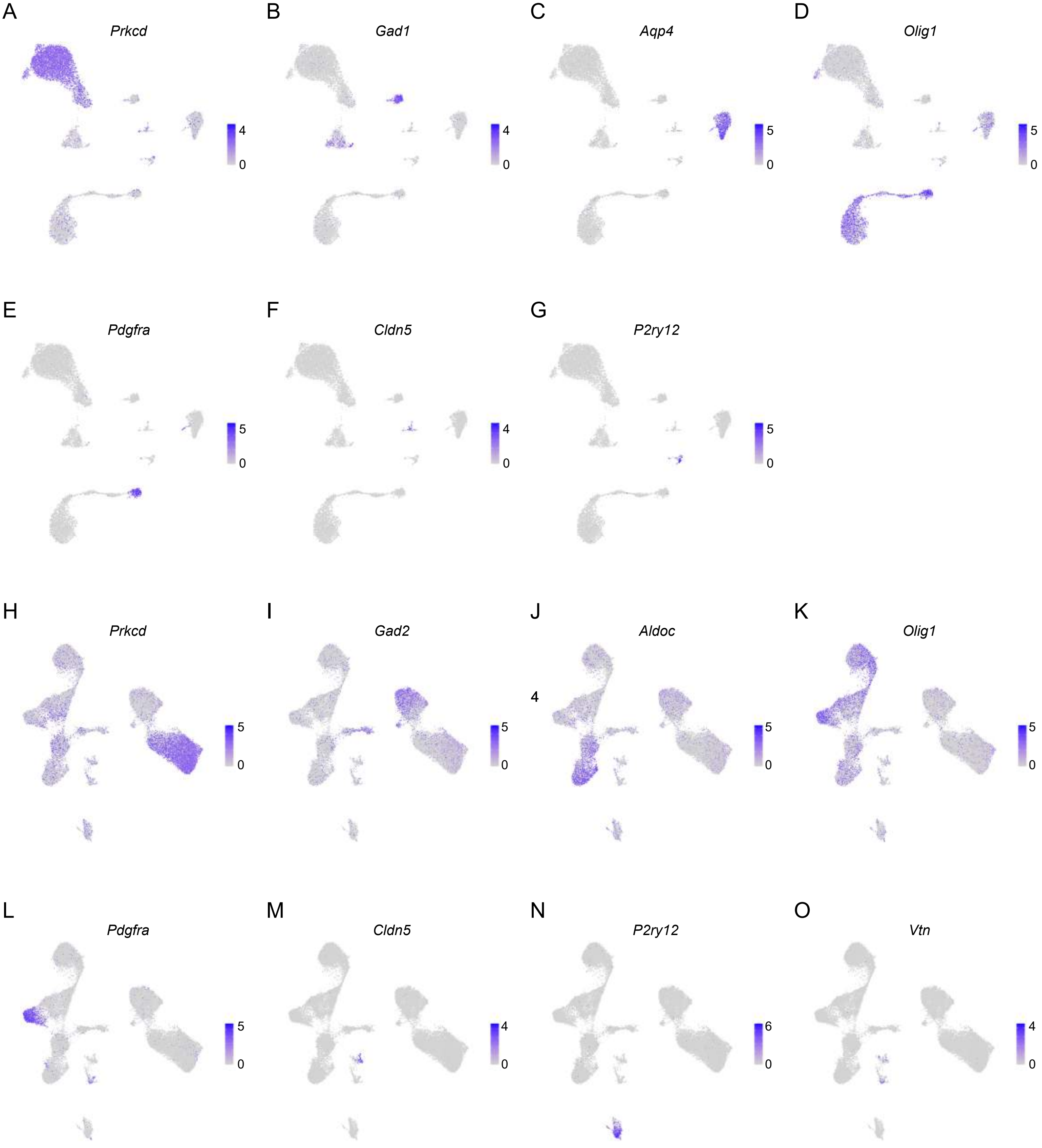


**Figure S2 (Related to figure 1). Marker gene expression in Fn14 KO and TWEAK KO datasets.** (A) - (G) Feature plots of marker gene expression in Fn14 KO dataset (scale, log normalized transcripts per cell). (H) - (O) Feature plots of marker gene expression in TWEAK KO dataset. (scale, log normalized transcripts per cell). Cell type markers plotted: *Prkcd* (relay neurons), *Gad1/2* (inhibitory neurons), *Aqp4* (astrocytes), *Olig1* (oligodendrocytes), *Pdgfra* (oligodendrocyte precursor cells), *Cldn5* (endothelial cells), *P2ry12* (microglia), and *Vtn* (pericytes).

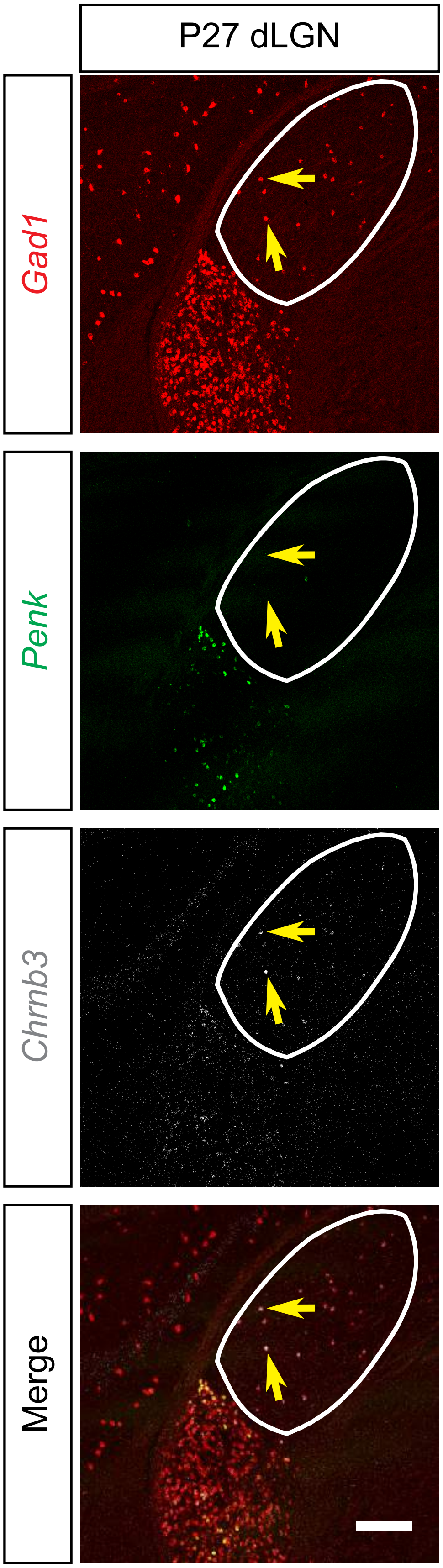


**Figure S3 (Related to figure 1)**. **Inhibitory neurons within and surrounding the dLGN cluster separately**. Low resolution confocal images of fluorescence *in situ* hybridizations (FISH, RNAscope) probing for the pan-inhibitory marker *Gad1* (red), dLGN-adjacent inhibitory neurons (*Penk;* Fn14 KO cluster 6, TWEAK KO cluster 2), and dLGN-resident inhibitory neurons (*Chrnb3*; Fn14 and TWEAK KO cluster 9). Scale bar, 200 µm.

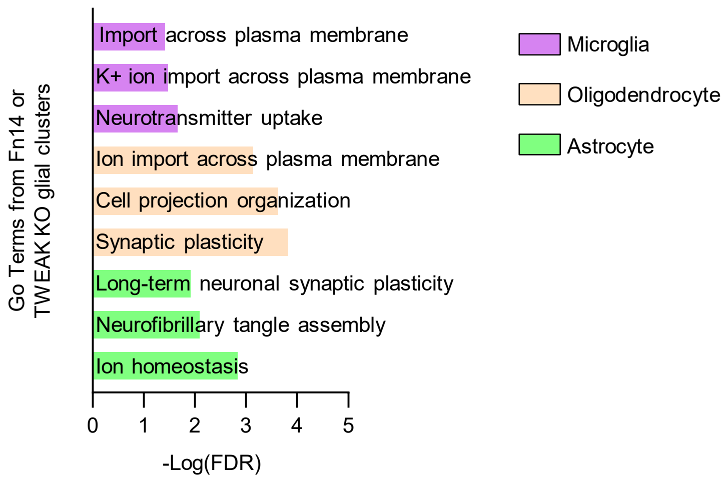


**Figure S4 (Related to figure 2)**. **Misregulation of genes in non-neuronal cells.** Select GO (gene ontology, PANTHER terms) categories enriched in significantly upregulated genes within TWEAK KO microglial clusters (purple), significantly downregulated genes within oligodendrocyte Fn14 KO clusters (orange), and significantly upregulated genes within astrocyte Fn14 KO clusters (green).

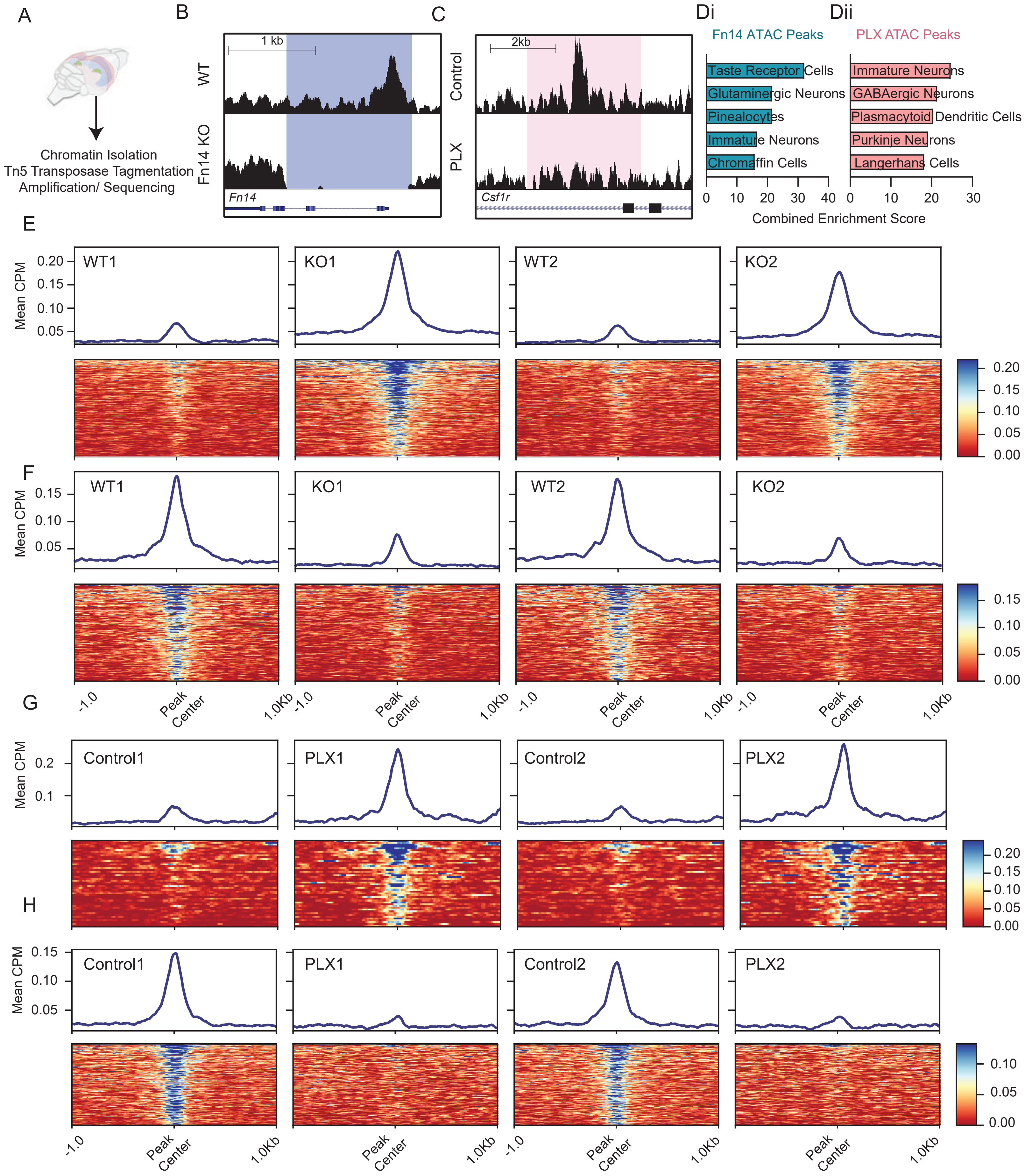


**Figure S5 (Related to figure 3).** Quality control and validation of Fn14 KO and PLX ATACseq datasets.

**Figure S5 (Related to figure 3). Quality control and validation of Fn14 KO and PLX ATACseq datasets.** (A) Schematic of a mouse brain depicting tissue taken from mice in Fn14 KO dataset (pink region, posterior brain) or PLX dataset (green region, dLGN). (B) Coding region of *Fn14* exemplifying loss of accessibility in the Fn14 KO tissue compared to WT tissue. (C) Coding region of *Csfr1* exemplifying loss of accessibility in the PLX tissue compared to control tissue. (Di)-(Dii) Cell type enrichment analysis based upon differentially accessible peaks in the Fn14 KO (Di) and PLX experiments (Dii). (E) Top: Mean counts per million (CPM) of peaks that gained accessibility in Fn14 KO tissues when compared to WT tissues. (Bottom) Heat maps of each peak in rank order of gained accessibility. (F) Top: Mean CPM of peaks that lost accessibility in Fn14 KO tissues. (Bottom) Heat maps of each peak in rank order of lost accessibility. (G) Top: Mean CPM of peaks that gained accessibility in PLX tissues when compared to Control tissues. (Bottom) Heat maps of each peak in rank order of gained accessibility. (H) Top: Mean CPM of peaks that lost accessibility in PLX tissues. (Bottom) Heat maps of each peak in rank order of lost accessibility. Each subpanel represents a separate bioreplicate.

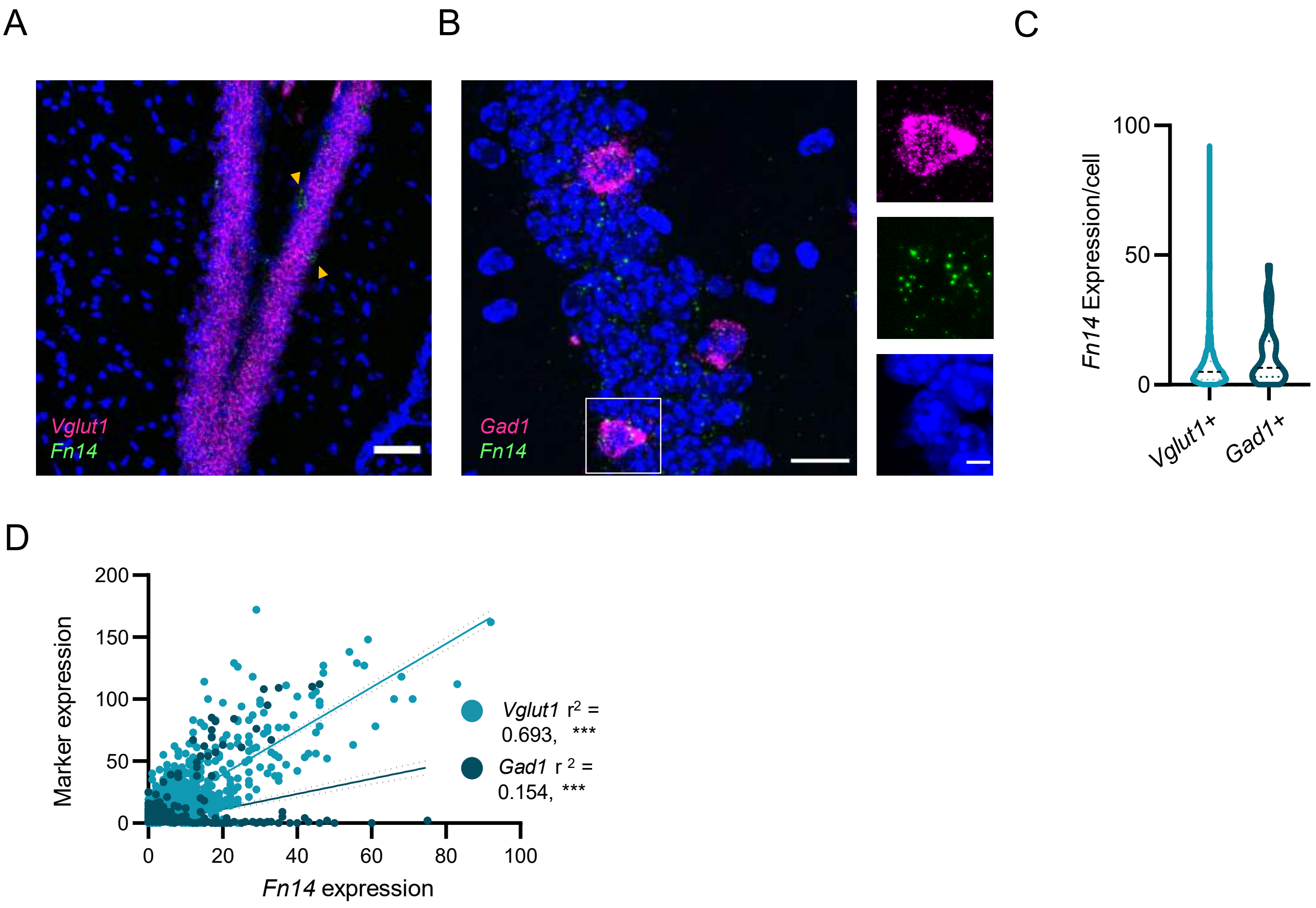


**Figure S6 (Related to figure 5). Rare *Gad1*+, *Fn14*+ cells in the hippocampus**. (A) confocal image displaying mRNA for *Fn14* (green) and the excitatory marker *Vglut1* (magenta). Triangles: *Fn14*+,*Vglut1*- cells. Scale bars, 40 μm. (B) higher magnification image of *Gad1*+ inhibitory cells expressing *Fn14*. *Fn14* (green), *Gad1* (magenta), and *Cldn5* (white). Scale bars, 20 and 5 μm for image and insert respectively. (C) Amount of *Fn14* in excitatory (*Vglut1*+) and inhibitory (*Gad1*+) neurons that express *Fn14* across ages. Violin plot of *Fn14* puncta/*Vglut1*+ (n = 1178 cells) or *Gad1*+ cells (n = 72 cells) ($p > 0.05$, Mann-Whitney). (D) Correlations between *Fn14* expression (x-axis) and marker gene expression (y-axis) reveals that *Fn14* expression is most closely correlated with *Vglut1* expression. Slope and $R^2$ values given in graph, and slopes were compared with linear regression analysis. Dotted lines represent each linear regressions' confidence (95% confidence intervals).

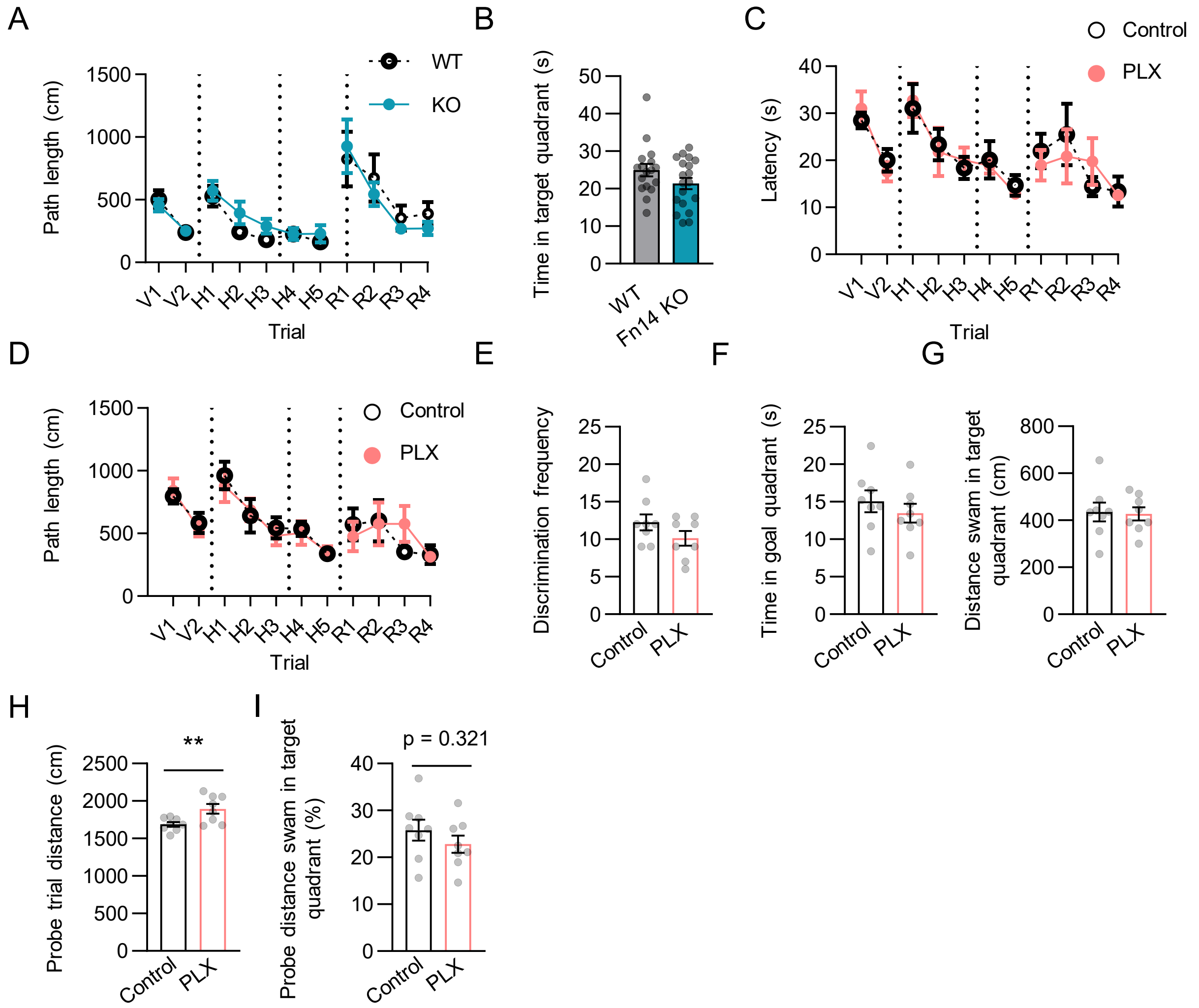


**Figure S7 (Related to figure 6). Learning and memory are relatively normal in mice lacking microglia.** (A) Path length swam by Fn14 WT and KO mice during the Morris Water Maze (MWM) test. (B) Time spent in goal quadrant by Fn14 WT and KO mice during MWM probe trials. (C)-(D) Latency to platform (C) and the path length (Distance; D) for control-fed and PLX-administered mice. Repeated measures ANOVA with Šídák's multiple comparisons test: Visible; treatment: $p > 0.05$, trial: $p < 0.05$, trial x treatment: $p > 0.05$, Hidden; treatment: $p > 0.05$, trial: $p < 0.05$, trial x treatment: $p > 0.05$, Reverse; treatment: $p > 0.05$, trial: $p < 0.05$, trial x treatment: $p > 0.05$. (E)-(G) Discrimination frequency of target quadrant (E), time in target quadrant (F), and distance swam in target quadrant (G) during probe trial of both control and PLX treated mice. $p > 0.05$ Student’s T test. (H) Total distance swam by control and PLX mice during probe trial. $p < 0.01$ Student’s T test. (I) Distance swam in target quadrant normalized to total distance swam during probe trail for control and PLX-treated mice (Student’s T-test, $p > 0.05$).

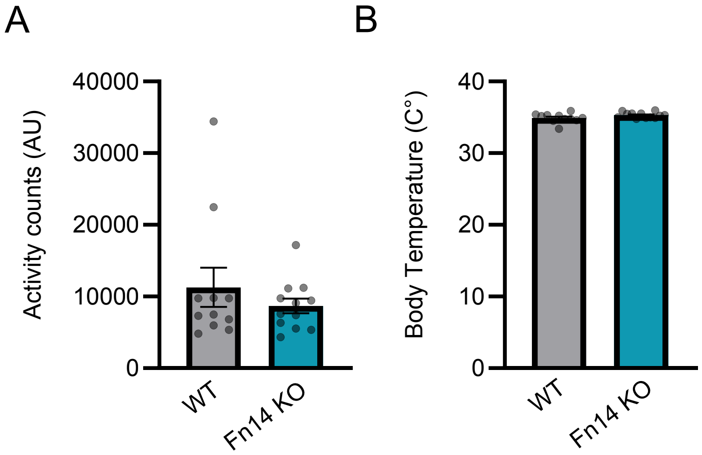


**Figure S8 (Related to figure 7). Fn14 KO mice exhibit normal activity and body temperature**. (A) Overall activity levels in Fn14 KO and WT mice as measured by automated detection by video recording during the EEG experiment. (B) Body temperatures were largely equivalent in Fn14 KO and WT mice. Two-tailed student's T test, $p > 0.05$.
